# Developmental disruption of the mitochondrial fission gene *drp-1* extends the longevity of *daf-2* insulin/IGF-1 receptor mutant

**DOI:** 10.1101/2024.02.07.579360

**Authors:** Annika Traa, Jeremy M. Van Raamsdonk

## Abstract

The dynamic nature of the mitochondrial network is regulated by mitochondrial fission and fusion, allowing for re-organization of mitochondria to adapt to the cell’s ever-changing needs. As organisms age, mitochondrial fission and fusion become dysregulated and mitochondrial networks become increasingly fragmented. Modulation of mitochondrial dynamics has been shown to affect longevity in fungi, yeast, *Drosophila* and *C. elegans*. While disruption of the mitochondrial fission gene *drp-1* only mildly increases wild-type lifespan, it drastically increases the already long lifespan of *daf-2* insulin/IGF-1 signaling (IIS) mutants. In this work, we determined the conditions required for *drp-1* disruption to extend *daf-2* longevity and explored the molecular mechanisms involved. We found that knockdown of *drp-1* during development is sufficient to extend *daf-2* lifespan, while tissue-specific knockdown of *drp-1* in neurons, intestine or muscle failed to increase *daf-2* longevity. Disruption of other genes involved in mitochondrial fission also increased *daf-2* lifespan as did treatment with a number of different RNAi clones that decrease mitochondrial fragmentation. In exploring potential mechanisms involved, we found that deletion of *drp-1* increases resistance to chronic stresses and slows physiologic rates in *daf-2* worms. In addition, we found that disruption of *drp-1* increased mitochondrial and peroxisomal connectedness in *daf-2* worms, increased oxidative phosphorylation and ATP levels, and increased mitophagy in *daf-2* worms, but did not affect their ROS levels or mitochondrial membrane potential. Overall, this work defined the conditions under which *drp-1* disruption increases *daf-2* lifespan and has identified multiple changes in *daf-2;drp-1* mutants that may contribute to their lifespan extension.

## Introduction

Mitochondria are important contributors to organismal health. In addition to being the primary producer of cellular energy, mitochondria also play key roles in apoptosis, calcium regulation, redox homeostasis and inter-organelle communication [1]. In aged organisms, mitochondrial function as well as mitochondrial morphology become dysregulated [2]. Though the mechanisms by which mitochondria influence longevity are not completely understood, defects in mitochondrial function contribute to multiple age-related metabolic [3–9], cardiovascular [10, 11] and neurodegenerative diseases [12–17].

Mitochondria form an interconnected network within the cell where fusion allows individual mitochondrion to join mitochondrial networks while fission allows mitochondrion to separate from each other. Regulation of mitochondrial fission and fusion allows mitochondria to dynamically respond to environmental conditions [18]. Mitochondrial fission facilitates the clearance of dysfunctional mitochondria as well as the generation of new mitochondria [19–21]. Mitochondrial fusion facilitates the complementation of mitochondrial components for optimal mitochondrial function [19, 22–24].

As organisms age, their mitochondrial networks become increasingly fragmented and lose the ability to switch between fused and fragmented networks [25–27]. Neurons from individuals with neurodegeneration have highly fragmented mitochondrial networks [28–31]. Furthermore, dysregulated expression of fission and fusion proteins occurs in both aging and age-related disease [2, 16, 32]. Notably, increased mitochondrial fusion has been associated with increased longevity. Healthy human centenarians have highly connected mitochondrial networks compared to 27- and 75-year-old individuals [33]. Additionally, in *C. elegans*, increased mitochondrial fusion is required for the longevity of multiple long-lived mutants [34–36] and overexpression of mitochondrial fusion genes is sufficient to extend longevity and enhance resistance to exogenous stressors (Traa et al., in revision for *Aging Cell*).

In *C. elegans,* the mammalian Opa1 homolog, EAT-3, fuses the inner mitochondrial membrane and the mammalian mitofusin homolog, FZO-1, fuses the outer mitochondrial membrane [37–39]. The protein responsible for the scission of both mitochondrial membranes in *C. elegans*, is DRP-1, homolog to the mammalian Drp1 [32, 40]. DRP-1 is recruited by FIS-1, FIS-2, MFF-1 and MFF-2 to the mitochondrial constriction site [41–45], where ER tubules wrap around mitochondria to begin mitochondrial membrane constriction [46, 47]. Oligomerization of DRP-1 into a ring structure occurs at the constriction site where DRP-1 hydrolyzes GTP for the energy needed to complete the scission of the inner and outer mitochondrial membrane [48, 49]. In addition to mitochondrial fission, DRP-1 also mediates peroxisomal fission and thus contributes to the regulation of peroxisomal network morphology [50].

Altering mitochondrial dynamics can affects an organism’s health and longevity. Disruption of mitochondrial fission increases lifespan in yeast and fungal models [51, 52]. In *C. elegans*, deletion of *drp-1* has little or no effect on lifespan in wild-type worms [35, 53] despite increasing resistance to specific exogenous stressors [25]. However, disruption of *drp-1* has been shown to affect longevity in other backgrounds. We found that disruption of *drp-1* increases lifespan and restores motility in a neuronal model of polyglutamine toxicity [54], but decreases lifespan in a model in which the toxic polyglutamine tract is expressed in muscle [55], suggesting that the beneficial effects of *drp-1* inhibition may be tissue-specific. Most notably, disruption of *drp-1* has been shown to extend the already long lifespan of *daf-2* insulin/IGF-1 receptor mutants and *age-1* phosphoinositide 3-kinase (PI3K) mutants [56].

The insulin/IGF-1 signaling (IIS) pathway is highly conserved among animals and links nutrient availability to organismal growth, metabolism and longevity [57]. The long lifespan of IIS pathway mutants, including *daf-2* and *age-1*, is dependent on the activation of the FOXO transcription factor DAF-16 [58]. DAF-16 upregulates the expression of pro-survival genes, such as chaperones and antioxidants, in response to stress, low nutrient availability, or when IIS is disrupted [59].

The mechanism by which *drp-1* extends *daf-2* lifespan is not known. Previous studies have indicated that *drp-1* deletion does not extend *daf-2* by enhancing DAF-16 activity, as disruption of *drp-1* does not increase DAF-16 activity in wild-type animals [25] or in *daf-2* mutants [56]. It has also been shown that the interaction between *drp-1* and the IIS pathway is not *daf-2-* specific as disruption of *drp-1* also extends the lifespan of *age-1* mutants [56]. Mutants of the IIS pathway have increased mitochondrial function, increased mitochondrial ROS production and increased induction of mitophagy, all of which may be affected by disrupting mitochondrial dynamics [60–62].

In this work, we define the conditions under which disruption of *drp-1* extends *daf-2* longevity and identify multiple factors that are altered by the loss of *drp-1* in *daf-2* mutants that may contribute to lifespan extension. We find that inhibition of *drp-1* during development is sufficient to increase *daf-2* lifespan but tissue specific inhibition of *drp-1* in the neurons, intestine or muscle fails to extend *daf-2* longevity. Additionally, decreasing mitochondrial fragmentation without disrupting *drp-1* is also sufficient to increase *daf-2* lifespan. We find that disruption of *drp-1* increases *daf-2* resistance to chronic stress, slows physiologic rates, increases mitochondrial function, increases mitophagy and enhances peroxisomal connectivity, all of which may contribute to the effect of *drp-1* on *daf-2* longevity.

## Methods

### Strains

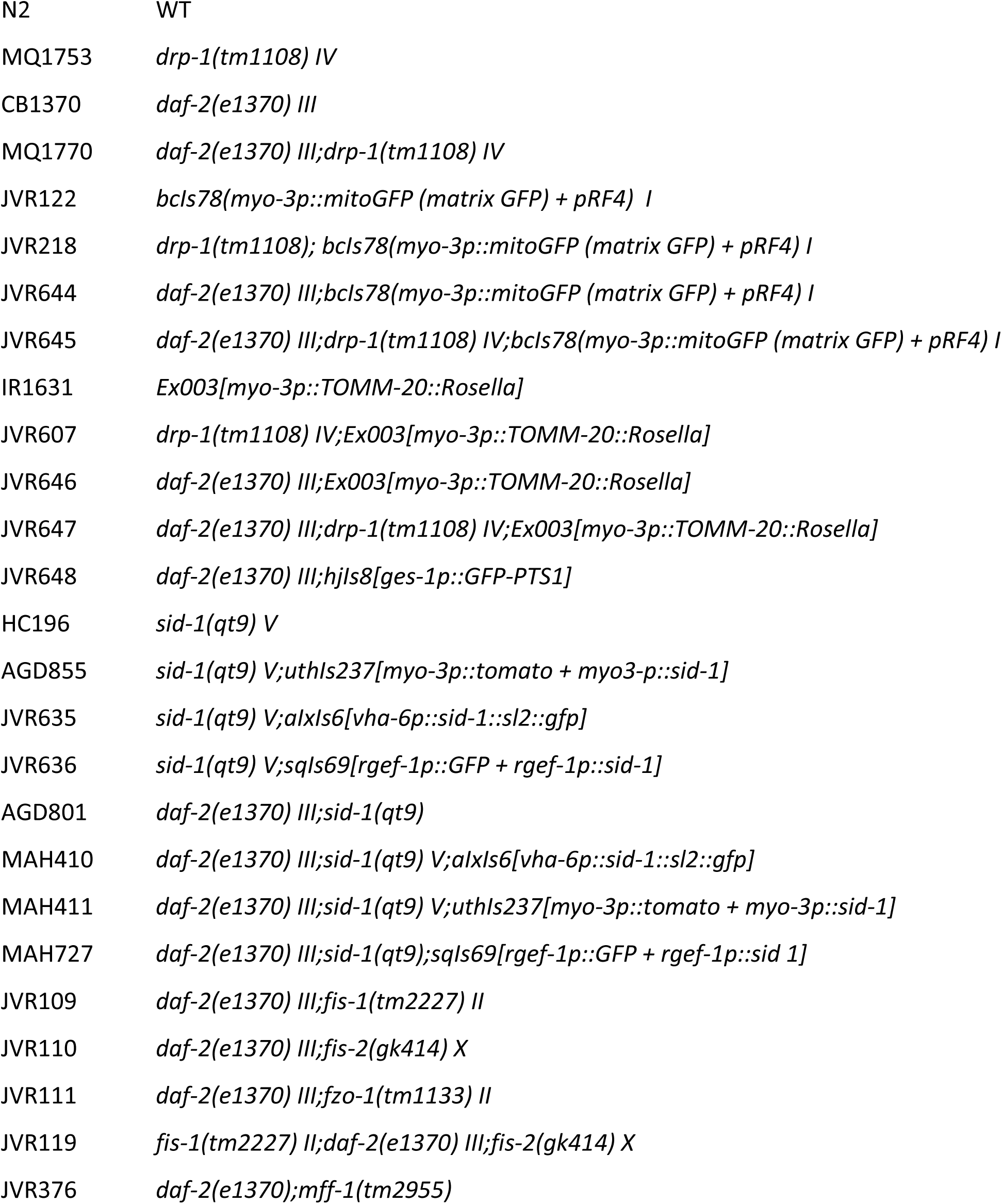
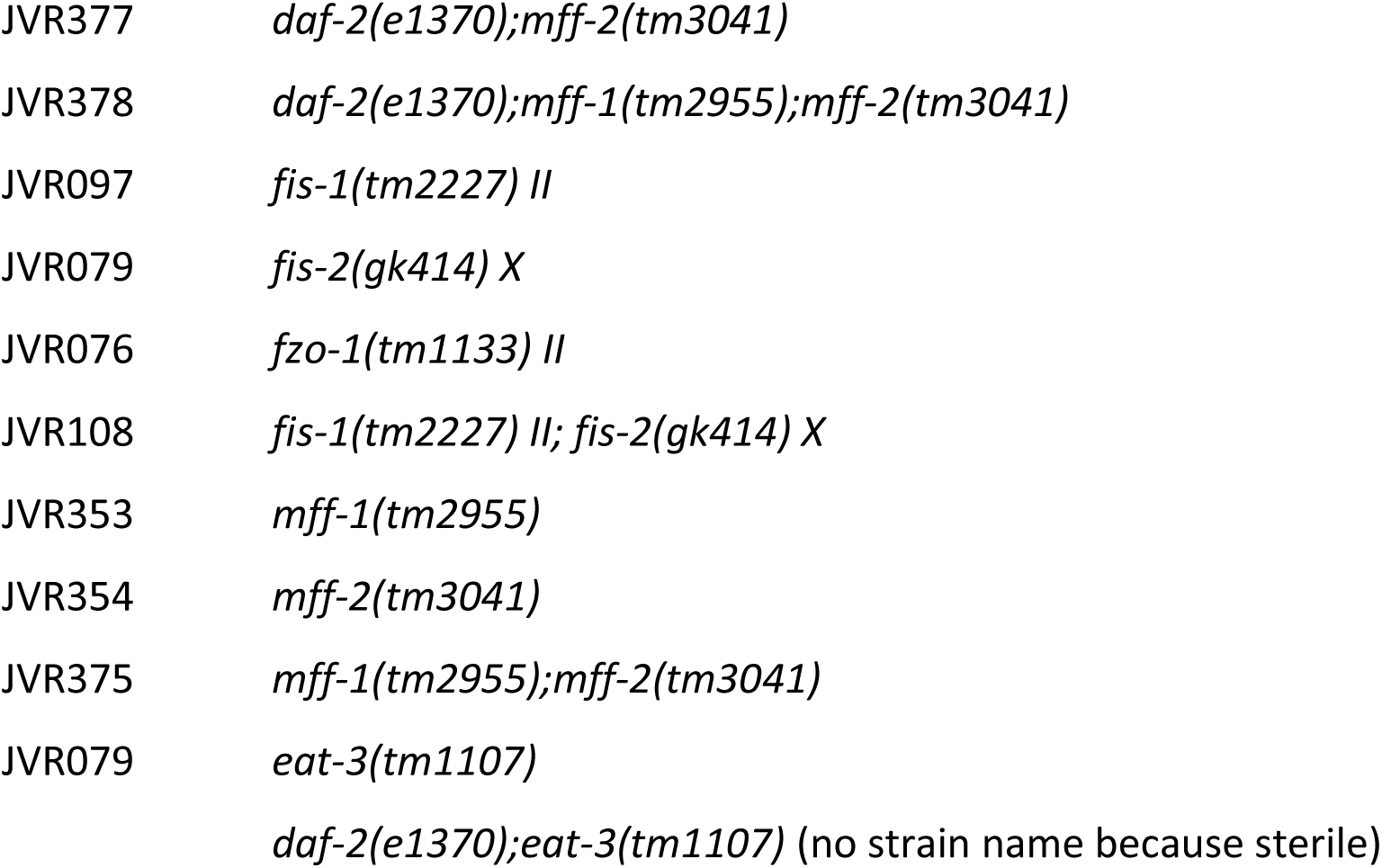

### Quantitative Real-Time RT-PCR

To perform quantitative RT-PCR, we first collected worms in M9 buffer and extracted RNA using Trizol as previously described [63, 64]. Using a High-Capacity cDNA Reverse Transcription kit (Applied Biosystems 4368814), the collected mRNA was then converted to cDNA. Quantitative PCR was performed using a PowerUp SYBR Green Master Mix (Applied Biosystems A25742) in a MicroAmp Optical 96-Well Reaction Plate (Applied Biosystems N8010560) and a Viia 7 Applied Biosystems qPCR machine. mRNA levels were calculated as the copy number of the gene of interest relative to the copy number of the endogenous control, *act-3*, then expressed as a percentage of wild-type. Primer sequences for each target gene are as follows: *drp-1* (L-GAGATGTCGCTATTATCGAACG, R-CTTTCGGCACACTATCCTG) *dcr-1* (L-ATTTTCGCGTCGTTAGCAGT, R-CGCATCATGTGGAAAATCAC).

### Confocal Imaging and Quantification

Mitochondrial morphology was imaged using worms that express mitochondrially-targeted GFP in the body wall muscle cells. In addition, we utilized a *rol-6* mutant background to facilitate imaging of the muscle cells [55]. The *rol-6* mutation results in animals moving in a twisting motion, allowing the sheaths of muscle cells to be facing the objective lens and thus more completely facilitates the imaging of mitochondrial networks within cells. Without the *rol-6* mutation, only the longitudinal edges of the muscle will often be visible, thus making it difficult to observe mitochondrial organization. Peroxisomal morphology was imaged using worms that express peroxisome targeted GFP in the intestine. Worms at day 1 or day 8 of adulthood were mounted on 2% agar pads and immobilized using 10 µM levamisole. Worms were imaged under a 40× objective lens on a Zeiss LSM 780 confocal microscope. Single plane images were collected for a total of twenty-four worms over three biological replicates for each strain. Imaging conditions were kept the same for all replicates and images. Quantification of mitochondrial or peroxisomal morphology was performed using ImageJ. Segmentation analysis was carried out using the SQUASSH (segmentation and quantification of subcellular shapes) plugin. Particle analysis was then used to measure the number, area, circularity, and maximum Feret’s diameter (an indicator of particle length) of the organelles.

The mtRosella mitophagy reporter was imaged as previously described in worms expressing the reporter in the body wall muscle [65]. The whole body of the worm was imaged under a 20x objective lens on a Zeiss LSM 780 confocal microscope. Quantification of dsRed and GFP fluorescence intensity was performed used ImageJ and representative images show both channels merged.

### Thrashing Rate

Thrashing rates were determined manually by transferring 20 age-synchronized worms onto an unseeded agar plate. One milliliter of M9 buffer was added and the number of body bends per 30 seconds was counted for 3 biological replicates of approximately 10 worms per strain.

### Brood Size

Brood size was determined by placing individual pre-fertile young adult animals onto NGM plates. Worms were transferred to fresh NGM plates daily until progeny production ceased. The resulting progeny was allowed to develop to the L4 stage before quantification. Three biological replicates of 5 animals each were completed.

### Oxygen Consumption Rate

Oxygen consumption measurements were taken using a Seahorse XFe96 analyzer [25, 66]. The night before the assay, probes were hydrated in 200 μL Seahorse calibrant at 37 degrees while the analyzer’s heater was turned off to allow the machine to cool. Day 1 and day 8 worms were collected in M9 buffer and washer three times before being pipetted into a Seahorse 96 well plate (Agilent Technologies Seahorse Flux Pack 103793-100). Others have previously determined that using between 5-25 worms per well is optimal [67]. Calibration was performed after 22 μL of FCCP and 24 μL sodium azide was loaded into the drug ports of the sensor cartridge. Measurements began within 30 minutes of worms being added to the wells. Basal oxygen consumption was measured 5 times before the first drug injection. FCCP-induced oxygen consumption was measured 9 times, then sodium-azide induced oxygen consumption was measured 4 times. Measurements were taken over the course of 2 minutes and before each measurement the contents of each well were mixed for an additional 2 minutes. Non-mitochondrial respiration (determined by sodium azide-induced oxygen consumption rate) was subtracted from basal respiration to calculate mitochondrial respiration.

### ATP Determination

Day 1 and day 8 adult worms were collected, washed 3 times and frozen in 50 μL of M9 buffer using liquid nitrogen. Samples were then immersed in boiling water for 15 minutes followed by ice for 5 minutes and finally spun down at 14,800g for 10 minutes at 4 ℃. Supernatants were diluted 10-fold before ATP measurements using a Molecular Probes ATP determination kit (A22066) and TECAN plate reader. Luminescence was normalized to protein content measured using a Pierce BCA protein determination kit.

### Heat Stress Assay

To measure resistance to heat stress, approximately 25 pre-fertile young adult worms were transferred to new NGM plates freshly seeded with OP50 bacteria and were incubated at 37℃. Starting at 12 hours, survival was measured every hour for a total of 18 hours of incubation. Three biological replicates were completed.

### Osmotic Stress Assay

To measure resistance to osmotic stress, approximately 25 pre-fertile young adult worms were transferred to NGM plates containing 700 mM NaCl and seeded with OP50 bacteria. Worms were kept at 20℃ for 24 hours before survival was scored. Five biological replicates were completed.

### Oxidative Stress Assays

Resistance to acute oxidative stress was measured by transferring approximately 25 pre-fertile young adult worms to 420 μM juglone plates seeded with OP50 bacteria. Worms were kept at 20℃ and survival was monitored every 2 hours for a total of 8 hours. Resistance to chronic oxidative stress was performed by transferring 30 pre-fertile young adult worms to freshly prepared plates containing 4 mM paraquat, 100 μM FUdR and seeded with concentrated OP50. Survival was monitored daily. Three biological replicates were completed for both assays.

### Anoxic Stress Assay

To measure resistance to anoxic stress, approximately 50 pre-fertile young adult worms were transferred to new NGM plates seeded with OP50 bacteria. To create a low-oxygen environment for the worms, we utilized Becton-Dickinson Bio-Bag Type A Environmental Chambers. Plates with young adult worms of each strain were placed in the Bio-Bags for 120 hours at 20℃, then removed from the bags and allowed to recover for 24 hours at 20℃ before survival was measured. Five biological replicates were completed.

### Bacterial Pathogen Stress Assay

We tested for nematode resistance to death by bacterial colonization of the intestine. The slow kill assay was performed as previously described [68, 69]. PA14 cultures were grown over night for a total of 16 hours and then seeded to the center of NGM agar plates containing 25 μM FUdR. Plates were left on the bench to dry overnight and were then incubated at 37℃ for 24 hours. Next the plates were left to adjust to the temperature at which the assay is conducted at 25℃ overnight. To begin the assay, approximately 50 age synchronized L4 worms were transferred to each plate. To monitor survival, deaths were scored twice a day until all worms had died. Three biological replicates were completed.

### Quantification of ROS levels

ROS levels were measured using dihydroethidium (DHE; ThermoFisher Scientific, D1168), as previously described [70, 71]. A 30 mM solution of DHE in DMSO was aliquoted and stored at −80 °C. When needed, 5 μl of DHE stock was diluted in 5 mL of PBS to create a 30 μM DHE solution. Age matched day 1 adult worms (approximately 50) were collected in PBS, transferred to a 1.5 mL centrifuge tube and washed 3 times in 1 mL PBS. To immerse worms in a final concentration of 15 μM DHE, 100 μL of PBS was left in the centrifuge tube after the final wash and 100 μL of 30 μM DHE was added to the tube. Centrifuge tubes were wrapped in tinfoil to protect from light and worms were incubated at room temperature, on a shaker for 1 h, then washed 3 times in PBS. For imaging, worms were mounted on a 2% agarose pad and immobilized with 10 mM levamisole. Worms were imaged with the 20 × objective using a Zeiss LSM 780 confocal microscope. A total of 60 worms were imaged over 3 biological replicates, per genotype. Image J was used to quantify the fluorescence intensity of ethidium-labelled ROS in the whole body of the worm. For each image (one worm per image), mean fluorescence intensity of the whole image and background intensity were measured. For each strain, autofluorescence was sampled from approximately 5-10 untreated worms and mean autofluorescence for the strain was determined. Fluorescence intensity of each worm was calculated by subtracting the image’s background fluorescence and the strain’s mean autofluorescence from the mean fluorescence of the image.

### Tetramethylrhodamine (TMRE) Staining

We used the potentiometric fluorescent indicator, TMRE, to measure mitochondrial membrane potential. TMRE was dissolved in DMSO to make a 50 mM stock solution. The TMRE stock solution was then diluted in M9 to make a 5 μM working solution. The working TMRE solution was then pipetted onto NGM plates seeded with OP50 and were left on a nutator for 30 minutes to allow the TMRE to spread evenly over the agar. Approximately 20 age matched L4 or day 7 worms were transferred to the TMRE plates which were then covered to protect from light and stored at 20 ℃ for 20 hours. To de-stain, worms were transferred to regular NGM plates seeded with OP50 and left at 20 ℃ for 4 hours. Worms were imaged with the 20 × objective using a Zeiss LSM 780 confocal microscope and ImageJ was used to quantify the fluorescence intensity of TMRE in the whole body of the worm.

### Lifespan Assay

Lifespan assays were completed at 20℃ and on NGM agar plates that contained FUdR to inhibit the development of progeny and limit internal hatching. We used a low concentration of 25 μM FUdR, which we have previously shown does not affect the longevity of wild-type worms [72]. For each lifespan assay, 40 pre-fertile young adult worms were transferred to 25 μM FUdR plates seeded with OP50 bacteria and were kept at 20℃. Four biological replicates were started on four subsequent days and all replicates were scored every other day to monitor survival until all worms died. Worms were excluded from the assay if they crawled off the agar and died on the side of the plate, had internal hatching of progeny or expulsion of internal organs. Raw lifespan data can be found in **Table S1**.

### RNAi

We used RNAi to inhibit the expression of specific genes. RNAi clones from the Ahringer library were streaked onto LB plates containing 50 μg/mL carbenicillin and 10 ug/ml Tetracycline. Individual colonies were then inoculated in 2YT media with 50 μg/mL carbenicillin and grown with aeration at 37 ℃ for 18 hours. For experiments testing the tissue and timing requirements for *drp-1* knockdown to extend lifespan, RNAi clones were sequence verified and were concentrated before being seeded onto NGM plates containing 3 mM IPTG and 50 μg/mL carbenicillin. Seeded RNAi plates were left on the bench for two days to induce dsRNA expression in the bacteria. When conducting lifespan assays, we used the L4 parental paradigm where RNAi knockdown is begun in the parental generation. Age matched L4 worms were transferred to RNAi plates for 24 hours after which they were transferred to another RNAi plate for 24 hours before being removed. The progeny from these worms were then transferred to the experimental RNAi plates containing 25 μM FUdR for lifespan assays. RNAi lifespan assays were conducted at 20 ℃ where deaths were scored every two days and a minimum of three biological replicates were completed.

### Statistical Analysis

A minimum of three biological replicates were completed for all assays. Where possible, the experimenter was blinded to the genotype during the course of the experiment, to ensure unbiased results. Statistical significance of differences between groups was determined by computing a t-test, a one-way ANOVA, a two-way ANOVA or a log-rank test using Graphpad Prism, as indicated in the figure legends. All error bars indicate the standard error of the mean.

## Results

### Disruption of *drp-1* extends lifespan and enhances resistance to chronic stress in *daf-2* mutants

To investigate the role of mitochondrial dynamics in the longevity of *daf-2* mutants, we disrupted the *drp-1* gene in *daf-2* mutants and measured their lifespan. While disruption of *drp-1* provides a slight but significant extension of lifespan in wild-type worms, disruption of *drp-1* drastically extends the lifespan of already long-lived *daf-2* mutants (**Fig. 1A**). In addition to having a significantly longer lifespan, *daf-2* is known to have an increased healthspan [73, 74] and increased resistance to stress [75–77] but reduced fecundity [78–80]. Thus, to evaluate how disruption of *drp-1* affects the physiology and biological resilience of *daf-2* worms, we measured motility (thrashing rate), fertility (brood size), and resistance to chronic oxidative stress (4 mM paraquat), acute oxidative stress (420 µM juglone), bacterial pathogen stress (*P. aeruginosa*, PA14), heat stress (37 ℃), osmotic stress (700 mM NaCl), and anoxic stress (72 hours, 24 hours recovery).

**Figure 1.**
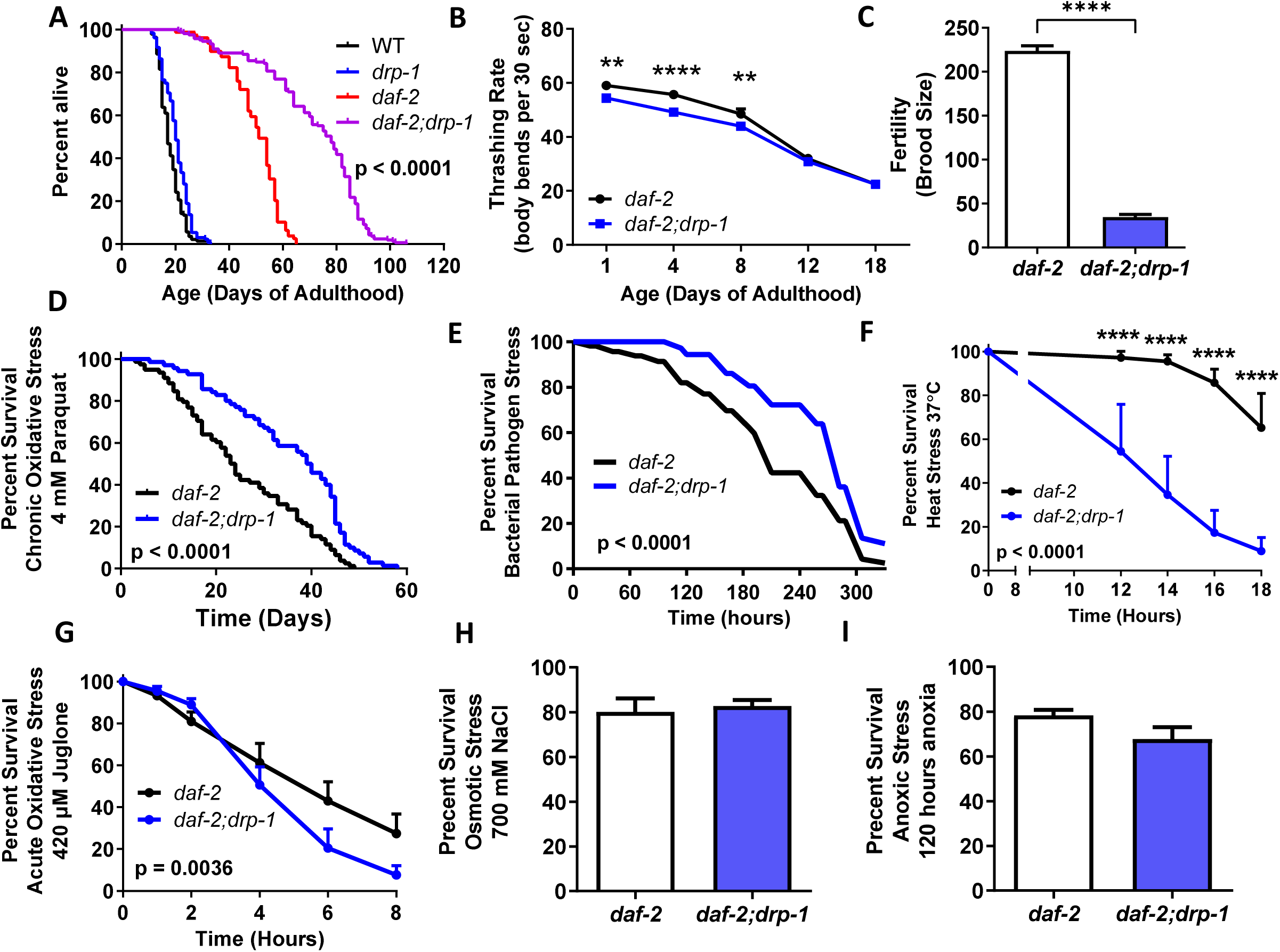
Disruption of *drp-1* extends the already long lifespan of *daf-2* mutants and increases resistance to bacterial pathogens and chronic oxidative stress. Deletion of *drp-1* markedly increases the lifespan of *daf-2* mutants **(A)**. Loss of *drp-1* reduces the rate of movement **(B)** and brood size **(C)** of *daf-2* worms. Deletion of *drp-1* increases *daf-2* resistance to chronic oxidative stress (**D**; 4 mm paraquat) and bacterial pathogen stress (**E**; *P. aeruginosa* strain PA14). In contrast, *daf-2;drp-1* mutants have increased sensitivity to heat stress (**F**) and acute oxidative stress (**G**). Deletion of *drp-1* does not affect resistance to osmotic stress (**H**) or anoxia (**I**) in *daf-2* worms. A minimum of three biological replicates were performed. Statistical significance was assessed using a log-rank test in panels A, D, E and G; a two-way ANOVA with Šidák’s multiple comparisons test for panel B; and a student’s t-test for panels C, H, and I. Error bars indicate SEM. **p<0.01, ****p<0.0001.

Disruption of *drp-1* slightly but significantly decreased the thrashing rate of *daf-2* animals at day 1, 4 and 8 of adulthood but had no effect at day 12 and 18 (**Fig. 1B**). Disruption of *drp-1* further decreased the brood size of *daf-2* worms such that a *daf-2;drp-1* double mutant produces few viable eggs (**Fig. 1C**). The *drp-1* deletion also slowed the development rate of *daf-2* worms.

We and others have shown that *daf-2* mutants have increased resistance to exogenous stressors [81, 82]. Disruption of *drp-1* in *daf-2* mutants further increased resistance to chronic stressors such as chronic oxidative stress (**Fig. 1D**) and bacterial pathogen stress (**Fig. 1E**). In contrast, disruption of *drp-1* decreased *daf-2* resistance to heat stress (**Fig. 1F**) and acute oxidative stress (**Fig. 1G**), but had no effect on resistance to osmotic stress (**Fig. 1H**) or anoxic stress (**Fig. 1I**). Combined these results show that the *drp-1* deletion slows physiologic rates and enhances resistance to specific stressors in *daf-2* worms, both of which have been associated with increased lifespan.

### Disruption of *drp-1* increases the connectivity of the mitochondrial and peroxisomal network in *daf-2* mutants

Using a strain expressing mitochondrially-targeted GFP in muscle cells, we examined how disruption of *drp-1* affects mitochondrial network morphology at day 1 and day 8 of adulthood in *daf-2* and wild-type animals. Similarly, to determine if disruption of *drp-1* affects peroxisomal network morphology in in *daf-2* worms, we visualized peroxisomes using a strain expressing peroxisome-targeted GFP in the intestine.

At day 1 of adulthood, disruption of *drp-1* produced a hyperfused mitochondrial network morphology in *daf-2* mutants, where rather than being organized into parallel elongated tubules, larger tubules and aggregated mitochondria were interconnected by thin filaments (**Fig. 2A**). Quantification of mitochondrial morphology revealed that deletion of *drp-1* in *daf-2* worms resulted in a decrease in the number of mitochondria, an increase in the average area of a mitochondrion, an increase in mitochondrial circularity and a decrease in the length of mitochondria (**Fig. 2A**). At day 8 of adulthood, the mitochondrial network of *daf-2;drp-1* mutants continues to appear more connected compared to the tubular morphology of *daf-2* mitochondrial networks, but both show an increase in mitochondrial fragmentation compared to day 1 adult worms as indicated by an increase in the number of mitochondria (**Fig. 2B**).

**Figure 2.**
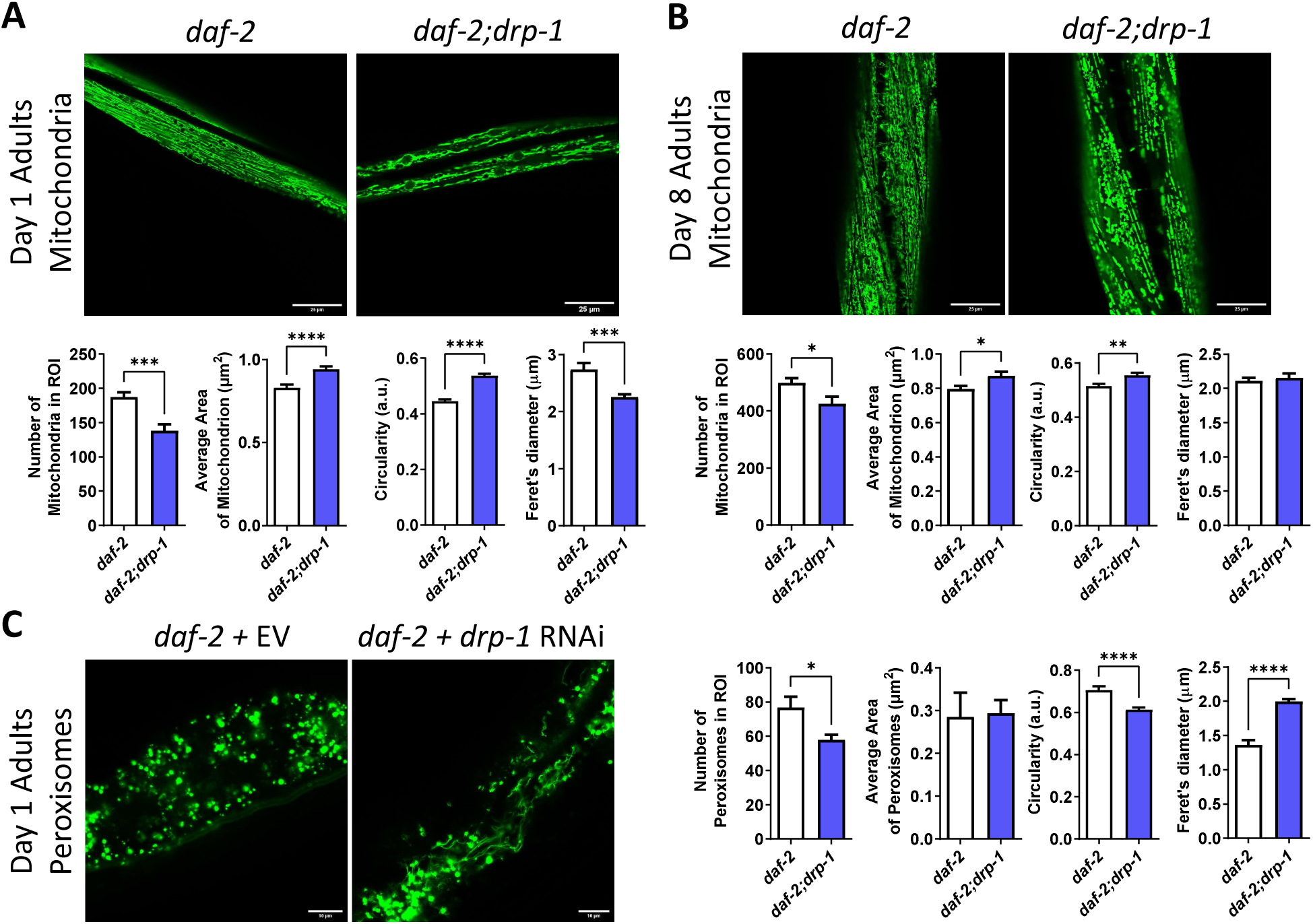
Disruption of *drp-1* increases mitochondrial and peroxisomal connectivity in *daf-2* mutants. **(A)** At day 1 of adulthood, deletion of *drp-1* decreases mitochondrial area, increases mitochondrial circularity and decreases mitochondrial length in *daf-2* worms. **(B)** At day 8 of adulthood, deletion of *drp-1* decreases mitochondrial number while increasing circularity in *daf-2* worms. **(C)** Decreasing *drp-1* levels with RNAi also affects peroxisome morphology in *daf-2* worms leading to decreased peroxisome numbers, decreased peroxisome circularity and increased peroxisome length. Three biological replicates were imaged. Statistical significance was assessed using a student’s t-test. Error bars indicate SEM. *p<0.05, **p<0.01, ***p<0.001, ****p<0.0001. Scale bar indicates 25 µm in panels A and B, and 10 µm in panel C.

Disruption of *drp-1* in wild-type animals also results in increased mitochondrial connectivity. At both day 1 (**Fig. S1A**) and day 8 (**Fig. S1B**) of adulthood, *drp-1* mutants exhibited a more fused mitochondrial network than wild-type worms as indicated by a decrease in the number of mitochondria, an increase in the average mitochondrial area, a decrease in the mitochondrial circularity and an increase in the average mitochondrial length.

Visualization of *daf-2* intestinal peroxisomes at day 1 of adulthood revealed that RNAi inhibition of *drp-1* increased peroxisomal network connectivity (**Fig. 2C**). In *daf-2* animals that were fed *drp-1* RNAi, peroxisomal tubules were visible resulting in a decrease in the number of peroxisomes, a decrease in peroxisomal circularity and an increase in peroxisomal length compared to *daf-2* animals that were fed empty vector bacteria. Thus, disruption of *drp-1* increases the connectivity of both the mitochondrial and peroxisomal networks in *daf-2* mutants.

### Inhibition of *drp-1* during development is sufficient to extend *daf-2* lifespan

In order to define the conditions under which *drp-1* disruption extends *daf-2* lifespan, we determined when during the animal’s life disruption of *drp-1* is required to extend *daf-2* lifespan. To do this, we fed *daf-2* and wild-type animals *drp-1* RNAi either during development only, during adulthood only, or during both development and adulthood and we measured their lifespans. Knockdown during adulthood was achieved by growing worms on empty vector bacteria and transferring worms to *drp-1* RNAi plates at day 1 of adulthood (**Fig. 3A**). Development only knockdown of *drp-1* was achieved by growing worms on *drp-1* RNAi until adulthood and then transferring these worms to RNAi targeting dicer (*dcr-1*), which is required for RNAi activity, in order to inhibit the knockdown of *drp-1* [83–85].

**Figure 3.**
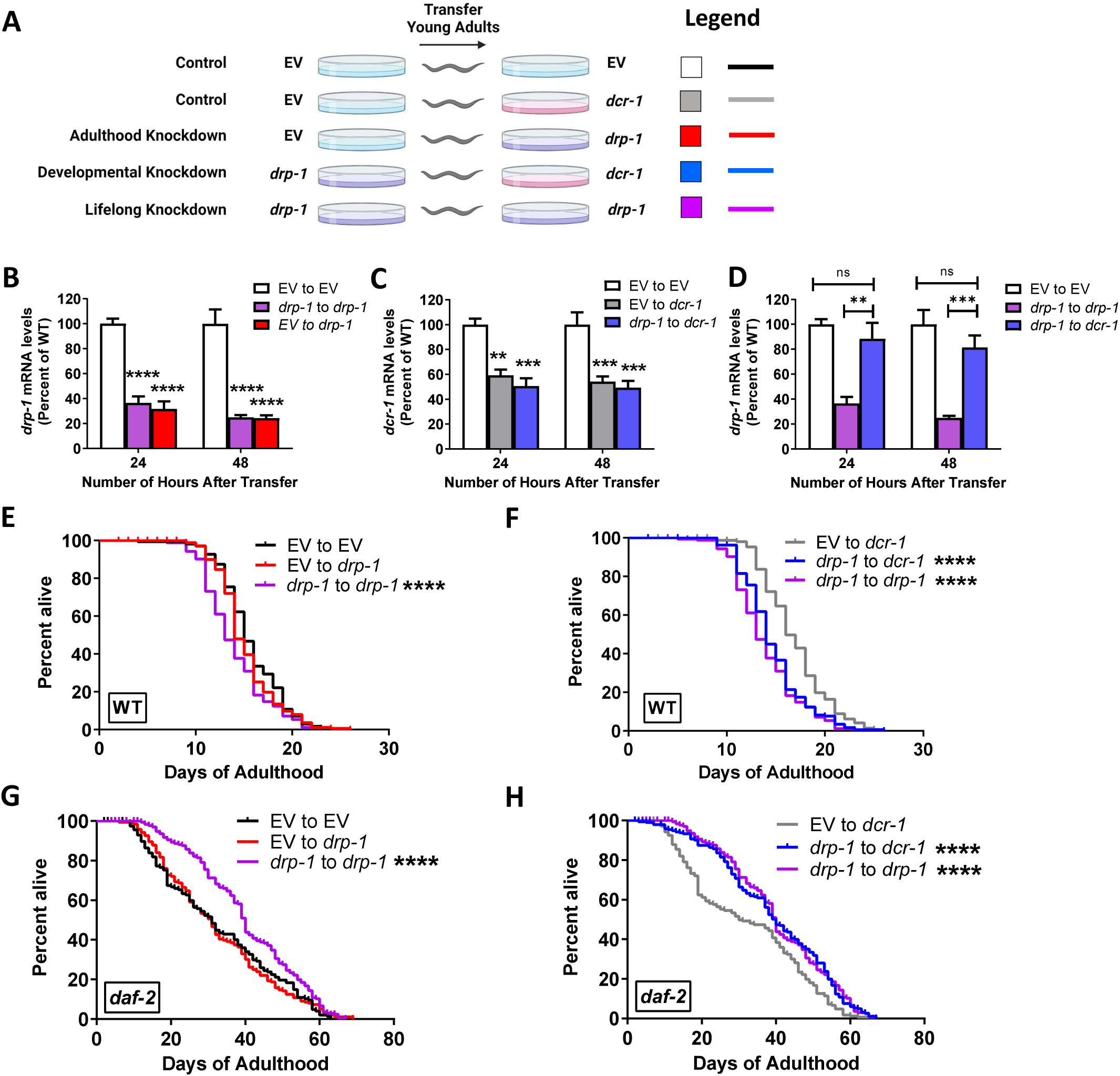
Inhibition of *drp-1* during development is sufficient to extend *daf-2* lifespan. **(A)** To determine when *drp-1* depletion acts to increase *daf-2* lifespan, *drp-1* levels were reduced during development only *(drp-1* to *dcr-1;* blue), during adulthood only (EV to *drp-1;* red) or during development and adulthood *(drp-1* to *drp-1;* purple) and compared to empty vector (EV) from development to adulthood (EV to EV; white bar or black line). **(B)** Measurement of *drp-1* levels by quantitative RT-PCR confirm that *drp-1* levels were decreased during adulthood by *drp-1* RNAi. **(C)** Worms transferred to *dcr-1* RNAi exhibit decreased levels of *dcr-1* mRNA. **(D)** Worms treated with *drp-1* RNAi during development and *dcr-1* RNAi during adulthood show a recovery of *drp-1* mRNA levels during adulthood. In wild-type worms, adulthood only knockdown of *drp-1* did not affect lifespan **(E)**, while development only *drp-1* RNAi decreased longevity **(F)**. In *daf-2* mutants, adult only *drp-1* RNAi did not affect longevity **(G)**, while *drp-1* knockdown during development significantly increased lifespan **(H)**. Three biological replicates were performed for panels B, C, and D; and four biological replicates were performed for panels E-H. Statistical significance was assessed using a one-way ANOVA with Dunnett’s multiple comparisons test in panels B, C, and D; and a log-rank test in panels E-H. Error bars indicate SEM. **p<0.01, ***p<0.001, ****p<0.0001.

Prior to beginning the lifespan experiments, we measured mRNA levels of *drp-1* and *dcr-1* to ensure that *drp-1* expression was being knocked down as expected. Adult only knockdown of *drp-1* significantly decreased the levels of *drp-1* mRNA within 1 day of transferring to *drp-1* RNAi (**Fig. 3B**). The level of knockdown achieved was equivalent to the level in worms with life long *drp-1* knockdown. For the development only paradigm, we confirmed that transferring to *dcr-1* RNAi decreased levels of *dcr-1* transcripts (**Fig. 3C**) and resulted in a recovery in the level of *drp-1* transcripts (**Fig. 3D**) one day after being transferred.

Wild-type animals with adulthood *drp-1* knockdown had no change in lifespan compared to those which were fed empty vector bacteria throughout their lives (**Fig. 3E**). However, wild-type animals with developmental or lifelong *drp-1* knockdown had shorter lifespans compared to those that were fed empty vector bacteria (**Fig. 3F**). In *daf-2* mutants, animals with adulthood *drp-1* knockdown did not live longer than animals with lifelong *drp-1* knockdown (**Fig. 3G**). However, *daf-2* mutants with developmental *drp-1* knockdown lived significantly longer than animals fed empty vector bacteria, such that their survival curve was not statistically different from that of animals with lifelong knockdown of *drp-1* (**Fig. 3H**). Thus, disruption of *drp-1* during development is sufficient to increase *daf-2* lifespan.

### Tissue specific inhibition of *drp-1* in the muscle, neurons or intestine is not sufficient to extend *daf-2* lifespan

To determine the extent to which inhibition of *drp-1* in specific tissues is sufficient to extend *daf-2* lifespan, we used RNAi to knockdown *drp-1* expression in the neurons, muscle and intestine of *daf-2* and wild-type animals. To achieve a tissue-specific knockdown of *drp-1*, we used *sid-1* mutants, which lack the dsRNA importer SID-1 in all of their tissues, and re-expressed *sid-1* from tissue specific promoters, resulting in tissue-specific RNAi sensitivity. The tissue-specific promoters that we used were *vha-6p* for the intestine, *myo-3p* for the muscle and *rgef-1p* for the neurons.

In order to confirm the tissue-specific RNAi sensitivity of each strain, we knocked down genes that act in each specific tissue to produce a clear phenotype. We knocked down the intestinal *elt-2* gene which causes L1 arrest (**Fig. 4A**), the hypodermal *bli-3* gene which causes cuticle blistering (**Fig. 4B**), the muscular *pat-4* gene which causes paralysis (**Fig. 4C**), and the neuronal *unc-70* gene which causes uncoordinated movement (**Fig. 4D**). We found that animals re-expressing *sid-1* from intestine-specific *vha-6p* were only sensitive to *elt-2* RNAi and animals re-expressing *sid-1* from the muscle-specific *myo-3p* were only sensitive to *pat-4* RNAi. Animals re-expressing *sid-1* from neuronal-specific *rgef-1p* were sensitive to *unc-70* RNAi and slightly sensitive to RNAi in the intestine.

**Figure 4.**
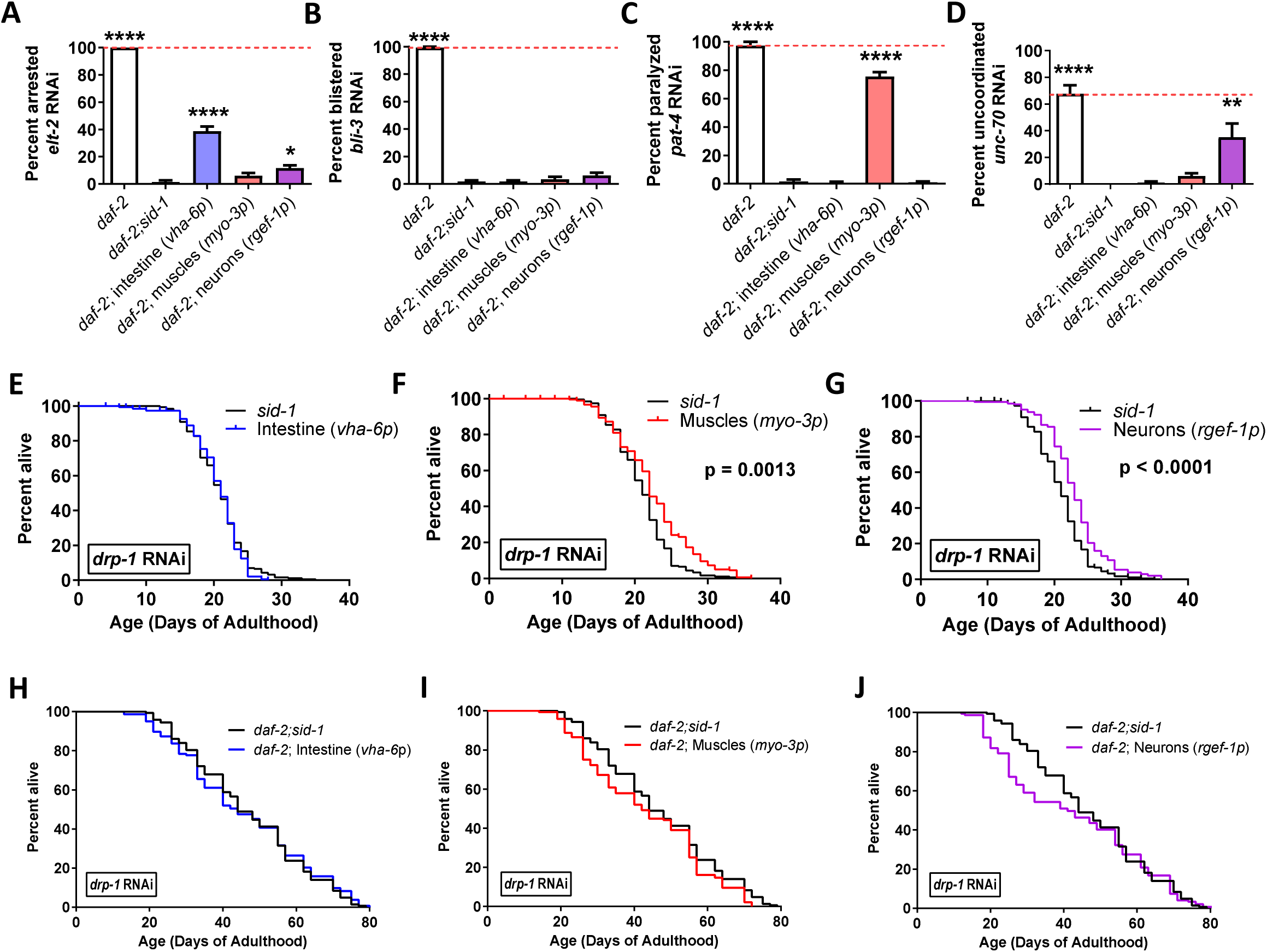
Inhibition of *drp-1* in individual tissues fails to extend *daf-2* lifespan. To identify the tissues in which decreasing the levels of *drp-1* acts to extend *daf-2* lifespan, tissue-specific RNAi was used to lower *drp-1* levels in different tissues of *daf-2* mutants and wild-type controls. To confirm that tissue-specific RNAi was working, all of the tissue-specific RNAi strains were treated with RNAi that should only produce a phenotype when knocked down in intestine (**A**; *elt-2* RNAi leading to arrestment and **B**; *bli-3* RNAi leading to blistering), body wall muscle (**C**; *pat-4* RNAi leading to paralysis) or neurons (**D**; *unc-70* RNAi leading to uncoordinated movement). In each case, treatment with RNAi caused a phenotype in *daf-2* worms and the corresponding tissue-specific RNAi strain but not *daf-2;sid-1* worms or any of the other untargeted tissue-specific RNAi strains, thereby indicating that the strains exhibit tissue specificity. In a wild-type background, knocking down *drp-1* in the intestine had no effect on lifespan (**E**), while knocking down *drp-1* in muscle (**F**) or neurons (**G**) resulted in a small increase in lifespan. In *daf-2* worms, knocking down *drp-1* in the intestine (**H**) body wall muscle (**I**) or neurons (**J**) had no effect on longevity. Three biological replicates were performed in panels A-D and four biological replicates were performed in panels E-L. Statistical significance was assessed using a one-way ANOVA with Dunnett’s multiple comparisons test in panels A-D and a log-rank test in panels E-L. In panels A-D, statistically significant differences from *daf-2;sid-1* are shown. Error bars indicate SEM. *p<0.05, **p<0.01, ***p<0.001, ****p<0.0001. A comparison of *drp-1* RNAi to empty vector can be found in Figure S2.

In considering whether inhibition of *drp-1* in a specific tissue is sufficient to extend lifespan, we compared the survival curve of animals with tissue-specific *drp-1* knockdown to two controls: animals with tissue-specific RNAi sensitivity that were grown on empty vector bacteria; and *sid-1* mutants which have whole-body resistance to RNAi. We considered a tissue-specific knockdown of *drp-1* to affect lifespan if its curve was statistically different from that of its empty vector control as well as the *sid-1* mutant control.

In wild-type animals, inhibition of *drp-1* in the intestine increased lifespan compared to the empty vector control (**Fig. S2A**) but not compared to *sid-1* mutants (**Fig. 4E**). Inhibition of *drp-1* in the muscle of wild-type animals did not affect lifespan compared to the empty vector control (**Fig. S2B**) but increased lifespan compared to *sid-1* mutants (**Fig. 4F**). Pan-neuronal inhibition of *drp-1* in wild-type animals increased lifespan compared to both the empty vector control (**Fig. S2C**) and *sid-1* mutants (**Fig. 4G**). Combined, this indicates that disruption of *drp-1* in neurons promotes longevity.

In *daf-2* animals, inhibition of *drp-1* in the intestine, muscle or neurons had no effect on lifespan compared to both the empty vector (**Fig. S2D-F**) and *sid-1* control (**Fig. 4H-J**). Overall, these results demonstrate that *drp-1* knockdown in intestine, muscle or neurons is not sufficient to extend *daf-2* lifespan.

### Disruption of multiple mitochondrial fission genes increases *daf-2* lifespan

To determine the extent to which *drp-1*’s role in mitochondrial fission contributes to lifespan extension in *daf-2;drp-1* double mutants, we evaluated whether disruption of other genes involved in mitochondrial fission could also increase longevity in *daf-2* worms. We found that disruption of *fis-1* or *fis-1* and *fis-2* together significantly increases *daf-2* lifespan, while a deletion of *fis-2* alone did not extend longevity (**Fig. 5A-C**). Similarly, disruption of *mff-1* and *mff-2* together extends *daf-2* lifespan, while single deletions in either gene have no effect (**Fig. 5D-F**). In contrast, disruption of *fis-1, fis-2, mff-1, mff-2* or combinations of these genes did not extend the lifespan of wild-type worms (**Fig. S3A-F**). Combined, these results demonstrate that disrupting other genes involved in mitochondrial fission can also increase the lifespan of *daf-2* worms.

**Figure 5.**
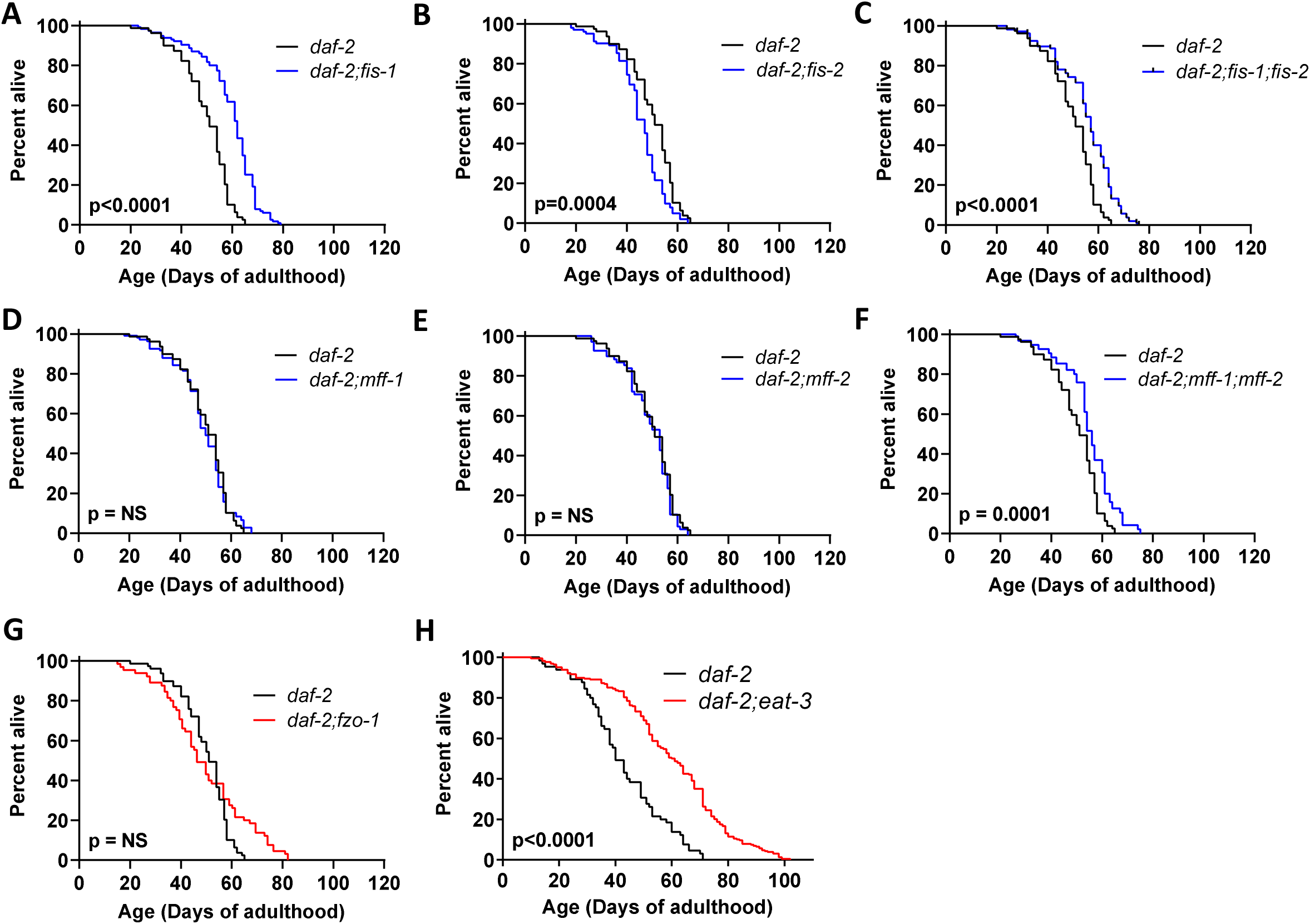
Disruption of mitochondrial fission genes increases *daf-2* lifespan. Deletion of the mitochondrial fission gene *fis-1* (**A**) or *fis-1* and *fis-2* together (**C**) increases *daf-2* lifespan, while disruption of *fis-2* results in a small decrease in *daf-2* lifespan (**B**). Disruption of the mitochondrial fission factor genes *mff-1* (**D**) or *mff-2* (**E**) do not affect *daf-2* lifespan individually but together result in a significant increase in *daf-2* longevity (**F**). In contrast to the ability of *drp-1* deletion to increase *daf-2* lifespan, disruption of *fzo-1* does not significantly affect *daf-2* lifespan (**G**). Disruption of *eat-3* markedly extends *daf-2* lifespan but also results in sterility (**H**). Three biological replicates were performed. Statistical significance was assessed using the log-rank test.

Since disruption of mitochondrial fission increases *daf-2* lifespan, we next evaluated whether disruption of mitochondrial fusion genes in *daf-2* worms would have the opposite effect. We found that disruption of *fzo-1* did not affect lifespan in *daf-2* mutants (**Fig. 5G**) or wild-type worms (**Fig. S3G**). While disruption of *eat-3* caused *daf-2* mutants to become sterile, the sterile *daf-2;eat-3* double mutants lived significantly longer than *daf-2* worms (**Fig. 5H**). Disruption of *eat-3* also extends lifespan in wild-type worms (**Fig. S3H**). Given that disruption of *eat-3* has multiple effects in *daf-2* worms, it is unclear whether the mechanism underlying lifespan extension is due to *eat-3*’s role in mitochondrial fusion, the effect of the *eat-3-*induced sterility on *daf-2* lifespan or the effect of dietary restriction on *daf-2* longevity (*eat* mutants have decreased feeding).

### Decreasing mitochondrial fragmentation without disrupting mitochondrial fission machinery can extend *daf-2* lifespan

As mitochondrial fission is important for the function of the cell, we tested whether decreasing mitochondrial fragmentation, without directly disrupting the mitochondrial fission machinery, could still extend *daf-2* lifespan. To do this, we treated *daf-2* worms with twenty-six RNAi clones that were previously shown to decreased mitochondrial fragmentation [86]. Of the twenty-six RNAi clones tested, ten RNAi clones significantly extended the lifespan of *daf-2* mutants (**Fig. 6A**), one decreased the lifespan of *daf-2* mutants and fifteen did not affect lifespan. RNAi clones that extended *daf-*2 lifespan include *sdha-2* (**Fig. 6B**), C34B2.8 (**Fig. 6C**), K02F3.2 (**Fig. 6D**), T10F2.2 (**Fig. 6E**), Y69F12A.b (**Fig. 6F**), Y69F12A.c (**Fig. 6G**), *timm-17B.1* (**Fig. 6H**), C33A12.1 (**Fig. 6I**), *cyp-35A1* (**Fig. 6H**) and *pgp-3* (**Fig. 6K**). These genes are involved in various pathways of metabolism, protein transport or oxidative phosphorylation and are not characterized as directly contributing to mitochondrial fission or fusion processes [86]. Combined, these results show that decreasing mitochondrial fragmentation, either through disruption of other mitochondrial fission genes (*fis-1, fis-2, mff-2, mff-2*) or genes that affect mitochondrial fragmentation but are not directly involved in mitochondrial fission, can be sufficient to increase *daf-2* lifespan without disrupting *drp-1*.

**Figure 6.**
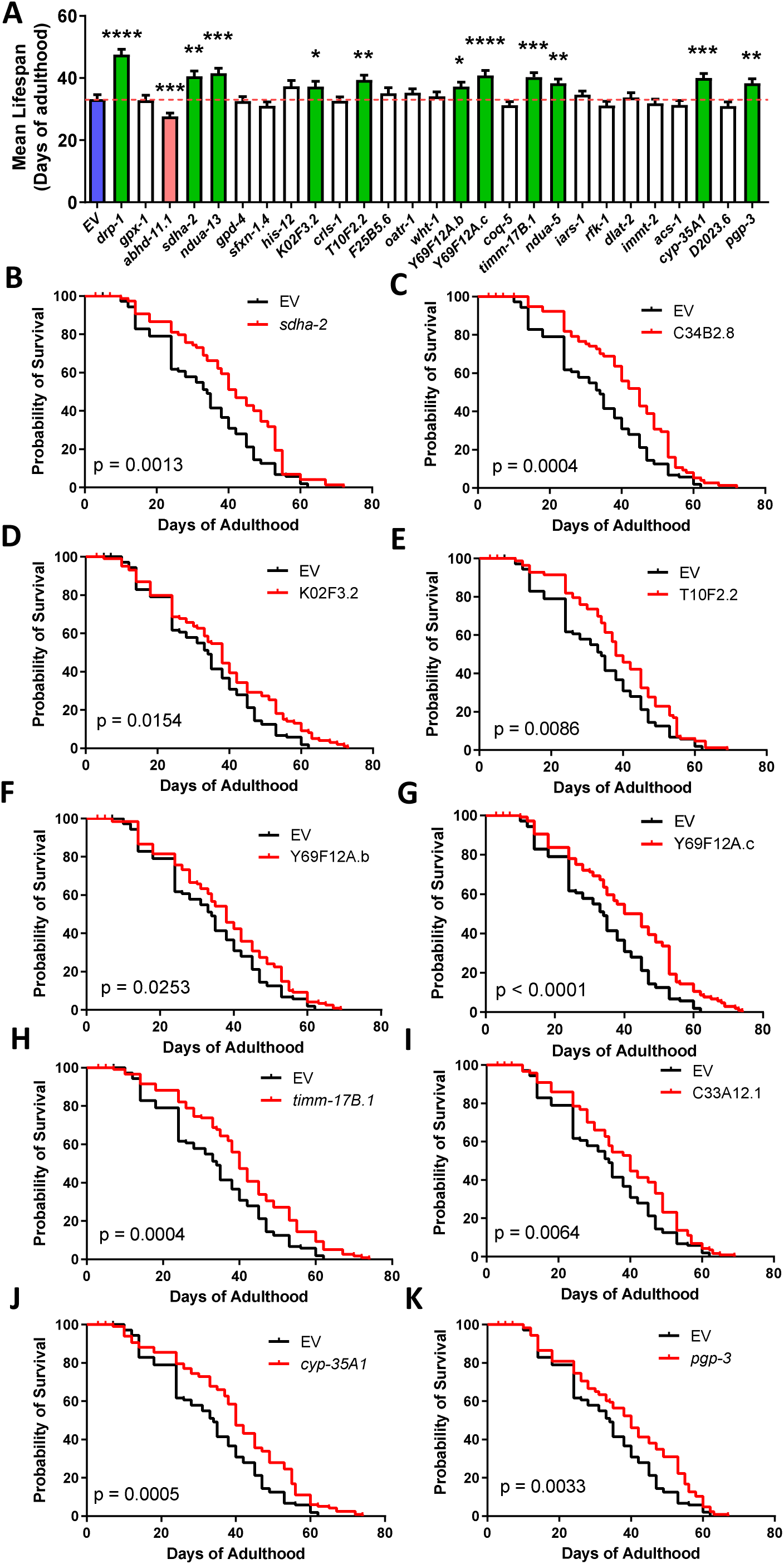
Decreasing mitochondrial fragmentation without disrupting *drp-1* can extend *daf-2* lifespan. To determine if decreasing mitochondrial fragmentation independently of *drp-1* could extend *daf-2* lifespan, *daf-2* worms were individually treated with 26 RNAi clones that were previously shown to decrease mitochondrial fragmentation in wild-type worms. Of these 26 RNAi clones, ten RNAi clones significantly increased the lifespan of *daf-2* mutants. This suggests that decreasing mitochondrial fragmentation may be sufficient to extend longevity in *daf-2* worms. RNAi clones that increased lifespan compared to control (blue) are shown in green. Three biological replicates were performed. Statistical significance was assessed using the log-rank test. Error bars indicate SEM. *p<0.05, **p<0.01, ***p<0.001, ****p<0.0001.

### Disruption of *drp-1* does not affect levels of ROS in *daf-2* mutants

*daf-2* worms and a number of other long-lived mutants have been shown to have increased levels of ROS, which contributes to their lifespan extension [62, 87–89]. As mitochondria are the primary site of ROS production in the cell, we asked whether disruption of *drp-1* might be altering ROS levels in *daf-2* worms and contributing to the extended lifespan of *daf-2;drp-1* double mutants. To determine whether ROS levels are altered by disruption of *drp-1* in either wild-type or *daf-2* mutants, we measured the fluorescence intensity of worms treated with the ROS-sensitive fluorescent dye dihydroethidium (DHE) at day 1 and day 8 of adulthood. While disruption of *drp-1* significantly increased the fluorescence intensity of DHE in wild-type animals, it did not further increase ROS levels in *daf-2* mutants (**Fig. 7A,B**). This suggests that altering ROS levels does not contribute to the effect of *drp-1* deletion on *daf-2* longevity.

**Figure 7.**
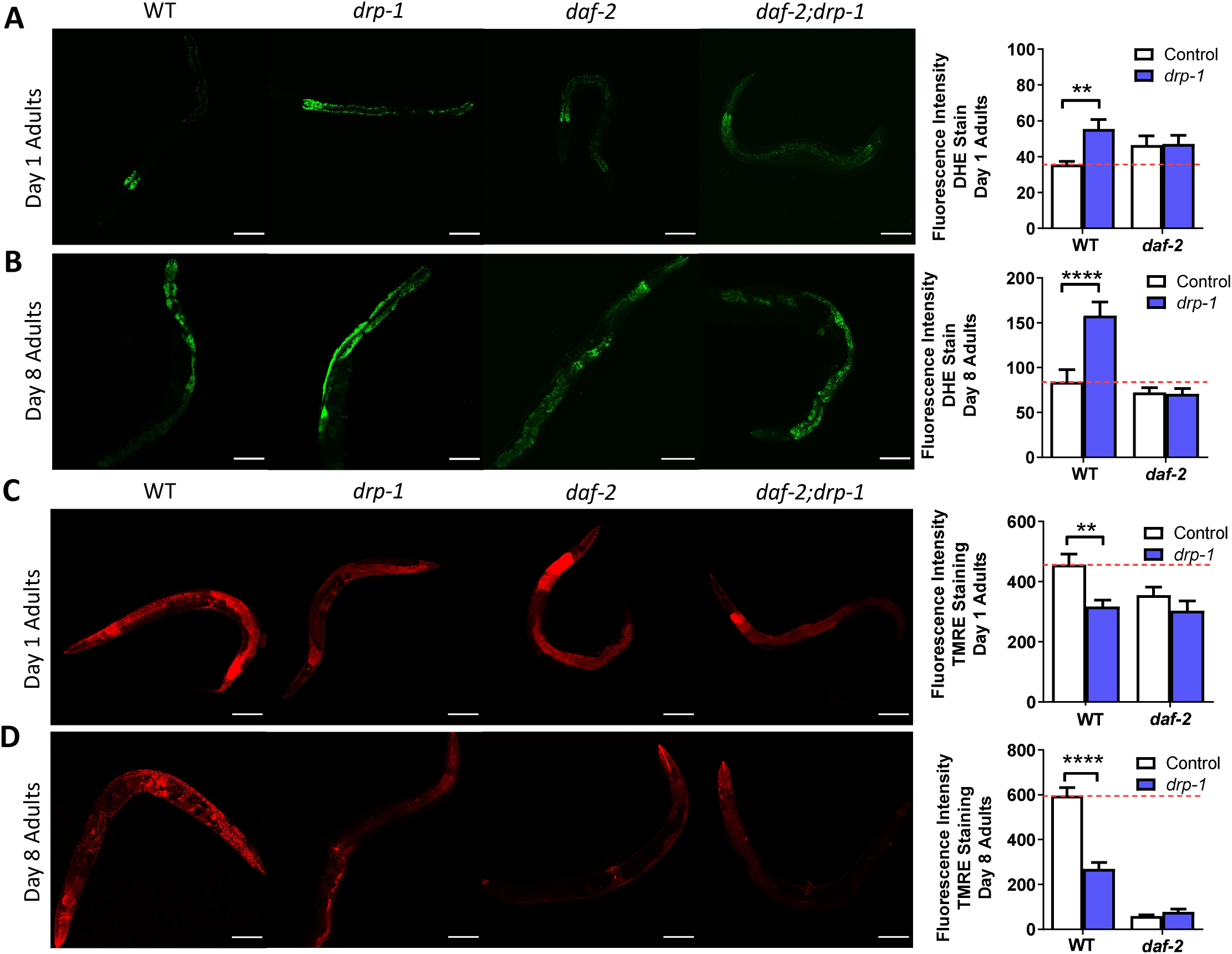
Disruption of *drp-1* increases ROS levels and decreases mitochondrial membrane potential in wild-type worms but not daf-2 mutants. ROS levels indicated by whole-worm dihydroethidium (DHE) staining are higher in *drp-1* worms compared to wild-type worms at day 1 of adulthood, while *daf-2;drp-1* worms do not have increased ROS levels compared to *daf-2* worms (**A**). Similarly, at day 8 of adulthood, ROS levels remain higher in *drp-1* worms compared to wild-type, while *daf-2;drp-1* worms continue to show ROS levels similar to *daf-2* (**B**). The mitochondrial membrane potential as indicated by whole-worm TMRE staining is decreased by disruption of *drp-1* in both wild-type worms at day 1 of adulthood (**C**). At day 8 of adulthood, disruption of *drp-1* decreases mitochondrial membrane potential in a wild-type background but not in a *daf-2* background (**D**). Three biological replicates were performed. Statistical significance was assessed using a two-way ANOVA with Šidák’s multiple comparisons test. Error bars indicate SEM. **p<0.01, ****p<0.0001. Scale bar indicates 100 µm.

### Disruption of *drp-1* does not affect mitochondrial membrane potential in *daf-2* mutants

Alterations in mitochondrial membrane potential have been associated with longevity. *daf-2* mutants and several other long-lived mutants were shown to have lower mitochondrial membrane potential than wild-type worms, though others have reported the opposite [60, 61, 90]. Mitochondrial membrane potential decreases with age [91] and preventing this decline is sufficient to increase lifespan [92]. Moreover, dietary restriction appears to increase lifespan at least partially through the preservation of the mitochondrial membrane potential [93]. Accordingly, we examined the effect of *drp-1* disruption on mitochondrial membrane potential in wild-type and *daf-2* worms to assess its potential contribution to lifespan extension.

To quantify mitochondrial membrane potential, we imaged day 1 and day 8 worms treated with the TMRE (tetramethylrhodamine ethyl ester), a fluorescent indicator whose uptake is dependant on mitochondrial membrane potential. At both time points, we found that *daf-2* worms have decreased mitochondrial membrane potential compared to wild-type animals (**Fig. 7C,D**). While disruption of *drp-1* decreased TMRE fluorescence in wild-type animals, it had no effect on TMRE fluorescence in *daf-2* mutants (**Fig. 7C,D**). This suggests that *drp-1* deletion does not increase *daf-2* lifespan through altering mitochondrial membrane potential.

### Disruption of *drp-1* increases mitochondrial function during young adulthood in *daf-2* mutants

Previous studies have reported alterations in mitochondrial function are associated with increased lifespan [61, 94–101]. To investigate whether disruption of *drp-1* affects mitochondrial function in *daf-2* worms, we measured oxygen consumption and ATP content at day 1 and day 8 of adulthood.

In day 1 adults, disruption of *drp-1* increases the oxygen consumption rate of *daf-2* mutants, but not of wild-type animals (**Fig. 8A**). Furthermore, in day 1 adults, disruption of *drp-1* increases ATP levels in *daf-2* mutants, but not in wild-type animals (**Fig. 8B**). In day 8 adults, disruption of *drp-1* does not affect the oxygen consumption rate (**Fig. 8C**) or ATP levels (**Fig. 8D**) of either wild-type or *daf-2* mutants. Combined these results show that disruption of *drp-1* further improves the mitochondrial function of *daf-2* mutants early in adulthood.

**Figure 8.**
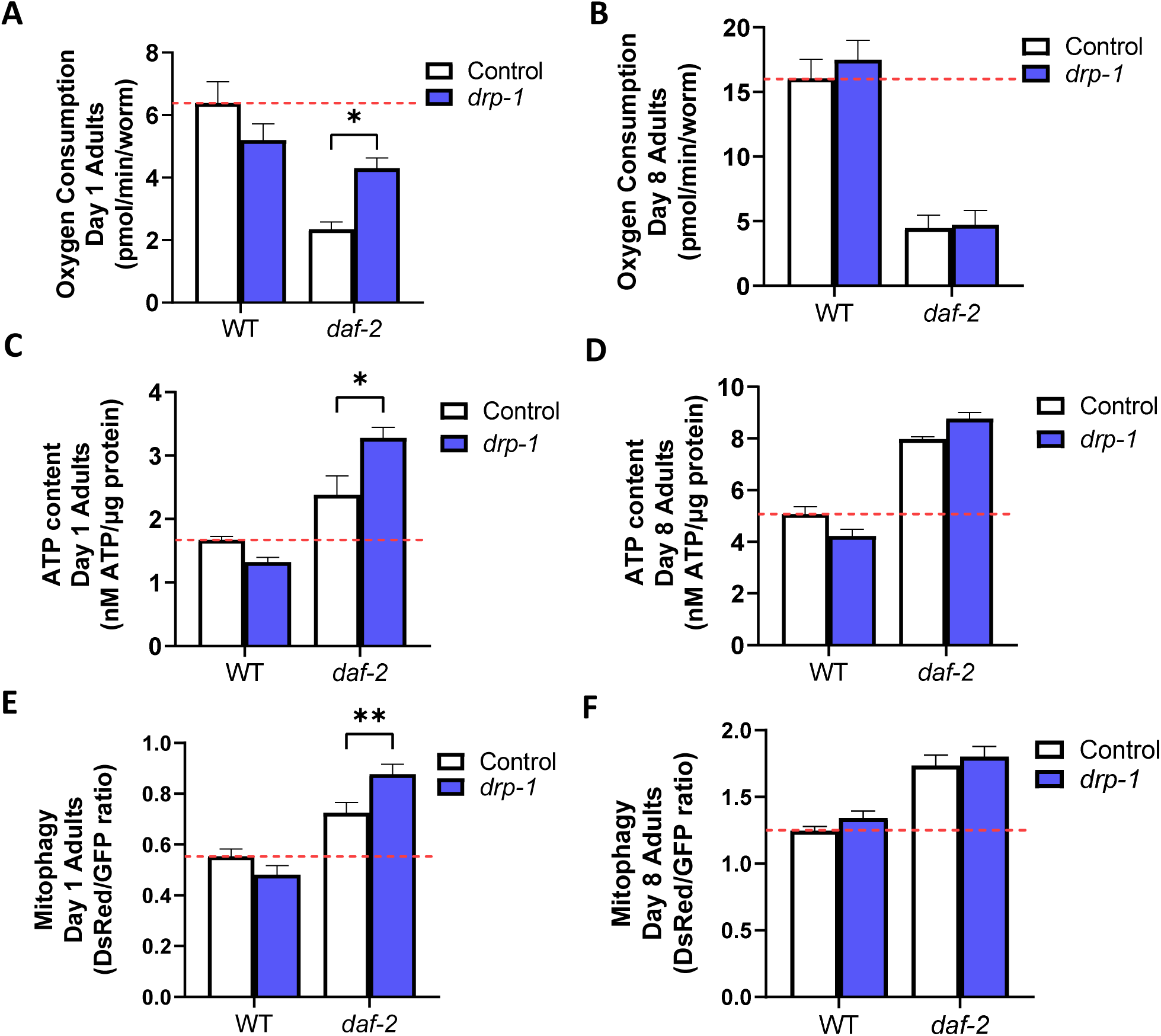
Disruption of *drp-1* increases mitochondrial function and mitophagy in *daf-2* mutants at day 1 of adulthood. While *drp-1* mutants consumed similar levels of oxygen as wild-type animals, *daf-2;drp-1* mutants consumed more oxygen than *daf-2* mutants at day 1 of adulthood (**A**). At day 8 of adulthood, disruption of *drp-1* does not affect the oxygen consumption (**B**). Similarly, *daf-2;drp-1* showed increased levels of ATP content compared to *daf-2* while *drp-1* mutants showed no change in ATP content compared to wild-type at day 1 (**C**), while at day 8 of adulthood, disruption of *drp-1* does not ATP levels (**D**). Quantification and comparison of the fluorescence intensity of both the pH-insensitive DsRed and pH-sensitive GFP fluorophores of the Rosella mitophagy reporter indicates that at day 1 of adulthood, disruption of *drp-1* induces mitophagy in *daf-2* mutants but not in wild-type worms (**E**). By contrast, at day 8 of adulthood, disruption of *drp-1* does not affect mitophagy in either wild-type or *daf-2* worms (**F**). Mitophagy induction is signified by an increase in the ratio of DsRed to GFP. Representative images of mtRosella staining can be found in **Figure S4**. Three biological replicates were performed. Statistical significance was assessed using a two-way ANOVA with Šidák’s multiple comparisons test. Error bars indicate SEM. *p<0.05, **p<0.01.

### Disruption of *drp-1* increases levels of mitophagy in *daf-2* mutants

Increased induction of mitophagy is observed in *daf-2* mutants [60] and has been shown to extend lifespan [102, 103]. As mitochondrial fission can facilitate mitophagy, we assessed whether disruption of *drp-1* affects levels of mitophagy in *daf-2* mutants. To do this, we generated *daf-2* animals expressing the mitochondrial-targeted Rosella (mtRosella) biosensor. mtRosella uses a pH-insensitive red fluorescent protein (dsRed) fused to a pH-sensitive green fluorescent protein (GFP) to monitor mitophagy levels in the body wall muscle cells of *C. elegans* [65]. After imaging the worms and quantifying the fluorescence intensity emitted by each protein, the dsRed to GFP ratio indicates levels of mitophagy. As expected, mitophagy levels were increased in *daf-2* mutants, compared to wild-type animals. Disruption of *drp-1* induces increased levels of mitophagy in *daf-2* mutants but not in wild-type animals at day 1 of adulthood (**Fig. 8E; Fig. S4**). At day 8 of adulthood, disruption of *drp-1* did not affect mitophagy levels in either wild-type or *daf-2* mutants (**Fig. 8F; Fig. S4**). Therefore, disruption of *drp-1* further increases mitophagy in *daf-2* mutants early in adulthood but has no effect on mitophagy in wild-type animals.

## Discussion

### Disruption of *drp-1* increases mitochondrial and peroxisomal network connectivity

In this work, we show that disruption of *drp-1* increases mitochondrial network connectivity in both *daf-2* and wild-type young adults. Later in adulthood, disruption of *drp-1* continues to generate increased mitochondrial network connectivity, but to a lesser extent. Increased mitochondrial network connectivity as a result of disrupting *drp-1* in wild-type animals has previously been documented [35, 39], although we and others have also previously observed no significant difference from wild-type mitochondrial morphology [25, 53, 104].

Using TMRE staining, the mitochondrial network morphology of *daf-2* mutants has previously been observed to be altered by disruption of *drp-1* at day 3 of adulthood [56]. The ratio of circularity to branching of the mitochondria was quantified and deemed to be higher when *drp-1* was disrupted in wild-type, *daf-2* and *age-1* animals. Here, we similarly observe an increase in mitochondrial circularity, however, we also consistently observe a decrease in the total number of mitochondria, and an increase in the average area of a mitochondrion, indicating that mitochondria are fusing together into larger aggregates. Increased mitochondrial fusion in *C. elegans* has previously been observed in several long-lived mutants, including *daf-2* [34]. Elongated mitochondrial tubules as well as lifespans were reported to be dependent on increased expression of the mitochondrial fusion protein EAT-3.

Given the ability of DRP-1 to regulate peroxisomal fission, we also considered the peroxisomal morphology of *daf-2* mutants and found that peroxisomes in the intestine are more interconnected when *drp-1* is disrupted. We observed an increase in filamentous structures, as visualized by GFP targeted to peroxisomes and these filaments appeared to connect circular structures together. Filamentous peroxisomal structures have previously been observed in *drp-1;fzo-1* worms, which require peroxisomal function for their increased longevity [35]. Thus, morphological changes in both the mitochondrial and peroxisomal network could both contribute to the longevity of *daf-2;drp-1* mutants.

### Disruption of *drp-1* further increases lifespan and stress resistance in *daf-2* mutants

In this work, we show that disruption of *drp-1* significantly enhances the already increased lifespan and stress resistance of *daf-2* mutants. The ability of *drp-1* disruption to increases *daf-2* and *age-1* lifespan was reported previously [56]. We also find that disruption of *drp-1* mildly increases wild-type lifespan. This is consistent with findings that disruption of *drp-1* can protect against age-associated pathologies in multiple models [54, 105–109], but contrasts with previous findings that *drp-1* mutants have a decreased lifespan [53] or have no change in lifespan in *C. elegans* [35].

To evaluate how disruption of *drp-1* affects the healthspan of *daf-2* mutants, we compared the thrashing rate of *daf-2* and *daf-2;drp-1* double mutants at multiple time points. We found that disruption of *drp-1* slightly but significantly decreases *daf-2* thrashing rate until day 8 of adulthood but does not affect thrashing rate later in adulthood. This data indicates that while disruption of *drp-1* may result in a slight decrease of *daf-2* motility early in life, it does not markedly affect *daf-2* healthspan. Disruption of *drp-1* does however significantly affect *daf-2* fertility, resulting in a drastic drop in the number of viable progeny produced by each worm. We and others have previously reported that *drp-1* mutants also have a significantly lower brood size compared to wild-type [25, 39]. This decrease in fertility could contribute to the ability of *drp-1* deletion to extend *daf-2* lifespan as many long-lived mutants have decreased fertility and complete ablation of the germline extends longevity [110, 111].

Though it was previously reported that disruption of *drp-1* decreases *age-1* resistance to chronic oxidative stress by paraquat exposure starting at day 3 of adulthood [56], we find that disruption of *drp-1* increases *daf-2* resistance to paraquat exposure starting at day 1 of adulthood. Additionally, we find that disruption of *drp-1* increases *daf-2* resistance to another chronic stress, bacterial pathogen stress by feeding worms *Pseudomonas aeruginosa*. We have not previously tested the resistance of *drp-1* mutants to bacterial pathogen stress, however, we have reported that *drp-1* mutants have increased resistance to paraquat exposure and increased activation of the SKN-1-mediated oxidative stress response pathway. We also showed that disruption of other mitochondrial fission genes such as *fis-1*, *fis-2, mff-1* and *mff-2* increases resistance to paraquat exposure [25].

Mitochondrial hyper-fusion has been observed in response to oxidative stress in a murine cell model, indicating that increased mitochondrial fusion may be beneficial in response to chronic oxidative stress, perhaps due to mitochondrial complementation mitigating the detrimental effects caused by small amounts of damage. By contrast, we find that disruption of *drp-1* decreases *daf-2* resistance to oxidative stress by juglone exposure and heat stress, both of which are acute stressors. Disruption of *drp-1* also decreases *age-1* resistance to heat stress [56]. In comparison, we have previously found that *drp-1* mutants have increased resistance to juglone exposure and decreased resistance to heat stress [25]. Thus, the ability of *drp-1* disruption to further increase *daf-2* resistance to chronic forms of stress may contribute to the ability of *drp-1* disruption to extend *daf-2* lifespan.

### Developmental disruption of *drp-1* increases *daf-2* lifespan

We find that developmental inhibition of *drp-1* is sufficient to increase *daf-2* lifespan. This finding is in agreement with our observation that disruption of *drp-1* affects *daf-2* mitochondrial morphology more significantly in early adulthood compared to later in life and indicates that the mechanism by which inhibition of *drp-1* extends *daf-2* lifespan takes place during development. Similarly, in long-lived mutants where altered mitochondrial function contributes principally to lifespan extension, genetic inhibition is required during development [85]. By contrast, *daf-2* inhibition is required during adulthood, and not during development, for lifespan extension to occur [84]. Thus, while mitochondrial dynamics and the IIS pathway converge to extend lifespan in *daf-2;drp-1* mutants, the mechanism by which disruption of *drp-1* extends *daf-2* lifespan occurs early in life, before disruption of *daf-2* contributes to lifespan extension.

### Tissue-specific disruption of *drp-1* does not increase *daf-2* lifespan

Our findings indicate that tissue-specific inhibition of *drp-1* in muscles, neurons or intestine fails to increase *daf-2* lifespan. This suggests that extension of *daf-2* lifespan either requires knockdown of *drp-1* in multiple tissues or that tissues not tested here, such as the germline, could be essential for lifespan extension mediated by disruption of *drp-1*. Inhibition of *daf-2* in the intestine or re-expression of *daf-16* in *daf-2;daf-16* mutants is sufficient to extend longevity [80, 112, 113], while re-expression of *daf-2* in the neurons or to a lesser extent the intestines decreases *daf-2* lifespan [114]. Combined with our findings here, this suggests that disruption of *drp-1* is not acting specifically in the same tissues as the *daf-2* mutation to increase lifespan.

In wild-type animals, we found that RNAi inhibition of *drp-1* in the neurons extends lifespan. This observation provides a plausible mechanism for why whole-body RNAi inhibition of *drp-1* decreases lifespan while deletion of *drp-1* increases lifespan. In the deletion mutant, *drp-1* is disrupted in all tissues including the neurons and the beneficial effect of *drp-1* disruption in neurons mediates the overall increase in lifespan. In wild-type worms treated with RNAi against *drp-1*, *drp-1* will be inefficiently knocked down in the neurons as RNAi is less effective in neuronal tissue [115–117]. Without the beneficial effect of disrupting *drp-1* in neurons, *drp-1* knockdown in the rest of the body has an overall detrimental effect on lifespan. In future studies, it will be interesting to more closely examine the mechanisms by which neuronal knockdown of *drp-1* extends lifespan.

### Decreasing mitochondrial fragmentation without disrupting *drp-1* can extend *daf-2* lifespan

We find that inhibiting genes known to be involved in mitochondrial fission as well as genes known to affect mitochondrial fragmentation is sufficient to extend *daf-2* lifespan without *drp-1* disruption. However, none of the targets tested were able to extend *daf-2* lifespan to the same magnitude as *drp-1*. This could be due to *drp-1* disruption having the strongest effect on mitochondrial fragmentation or that another activity of *drp-1* also contributes to its effect on *daf-2* lifespan.

Of the RNAi clones that decrease mitochondrial fragmentation and increased *daf-*2 lifespan, we previously reported that inhibition of *pgp-3* or *alh-12* is able to decrease mitochondrial network disorganization and improve movement in models of polyglutamine toxicity [55]. Furthermore, chemical inhibition of mitochondrial fission has previously been found to increase yeast lifespan during chronological aging [118], another example whereby decreasing mitochondrial fragmentation without inhibiting *drp-1* promotes lifespan extension. Limiting *drp-1* inhibition by identification of *drp-1-*alternative mechanisms to promote lifespan extension via decreased mitochondrial fragmentation may be useful given that *drp-1* disruption can be detrimental in mammals [119–121].

### Disruption of *drp-1* improves mitochondrial function in young *daf-2* adults

Mitochondrial dysfunction and an abnormally high or low mitochondrial membrane potential have both been associated with mitochondrial fragmentation and increased generation of mitochondrial ROS. Under normal physiological conditions, it is thought that increased mitochondrial network fusion promotes a mitochondrial membrane potential that is sufficiently elevated to enhance mitochondrial function without crossing the threshold into ROS overproduction [122]. Decreased membrane potential, or increased ROS levels may act as signals to induce mitochondrial fragmentation and mitophagy [123–125].

Though differing outcomes have been reported for the membrane potential of *daf-2* mutants [60, 61, 90], increased ATP production and increased ROS generation are consistently observed [61, 62, 88]. We find that disruption of *drp-1* in *daf-2* mutants has no effect on mitochondrial membrane potential or ROS levels but does increase mitochondrial function during early adulthood. Thus, redox homeostasis, which is maintained by DAF-16 and SKN-1 activity in *daf-2* mutants [126, 127], remains unaffected by the enhanced mitochondrial function mediated by *drp-1* disruption. Increased mitochondrial function in young *daf-2;drp-1* mutants, but not later in life, corresponds with our finding that disruption of *drp-1* during development is sufficient to extend *daf-2* lifespan.

We have previously observed that disruption of *drp-1* has no effect on mitochondrial function at day 1 or day 7 of adulthood [25]. Similarly, others have previously reported that *drp-1* mutants do not have altered ATP-coupled respiration, although increased basal respiration and an increased proton leak is reported [104]. While *drp-1* mutants would be expected to have increased mitochondrial function, membrane potential and mildly elevated ROS levels due to their increased mitochondrial network fusion, in this work, we again show that disruption of *drp-1* in wild-type animals does not affect mitochondrial function. Additionally, while we do see a significant increase in ROS levels in *drp-1* mutants, we also see a significant decrease in mitochondrial membrane potential. This elevated level of ROS and lowered mitochondrial membrane likely contributes to activation of the SKN-1-mediated oxidative stress response and may explain why disruption of *drp-1* is beneficial only under certain conditions. Additionally, the absence of enhanced mitochondrial function may help to explain why disruption of *drp-1* in wild-type animals does not extend lifespan as significantly as in *daf-2* mutants, where mitochondrial function is enhanced.

### Disruption of *drp-1* increases induction of mitophagy in young *daf-2* adults

Increased DAF-16 and SKN-1 activity leads to increased mitochondrial quality control by selective autophagy in *daf-2* mutants [128]. Inhibition of mitophagy partially decreases the lifespan of *daf-2* mutants, indicating that increased mitophagy is required to maintain *daf-2* longevity [60]. Although DRP-1 is required for selective clearance of dysfunctional mitochondria by recruitment of mitophagy machinery to specific membrane domains, DRP-1-independent mitophagy is possible [20, 129]. However, inhibition of fission machinery has been found to decrease mitophagy, resulting in the accumulation of oxidized mitochondrial proteins and reduced mitochondrial respiration, indicating that mitochondrial fission machinery plays an important role in mitochondrial quality control [21, 130].

Given that inhibiting DRP-1 has previously been found to affect mitophagy activity, we evaluated whether disruption of *drp-1* affects mitophagy in *daf-2* mutants. Unexpectedly, we find that disruption of *drp-1* further increases *daf-2* mitophagy but does not affect wild-type mitophagy levels. Furthermore, as with mitochondrial function, disruption of *drp-1* only enhances *daf-2* mitophagy in young adults and does not affect mitophagy later in life, corresponding with our finding that disruption of *drp-1* during development but not adulthood extends *daf-2* lifespan. While increased mitophagy in the absence of DRP-1, would be expected to be excessive and non-selective [20], the enhanced mitochondrial function and maintained ROS and membrane potential seen in *daf-2;drp-1* mutants suggest that the increased mitophagy remains beneficial.

## Conclusion

Overall, this work defines the conditions under which disruption of *drp-1* extends *daf-2* lifespan and identifies multiple putative mechanisms that contribute to lifespan extension. *drp-1* inhibition acts during development to extend *daf-2* lifespan in multiple tissues or tissues other than intestine, neurons and muscle. Disruption of *drp-1* increases mitochondrial and peroxisomal connectedness, decreases reproduction, enhances resistance to chronic stress, increases mitochondrial function and increases mitophagy in *daf-2* mutants, all of which may contribute to lifespan extension. Similar to *drp-1*, decreasing mitochondrial fragmentation by targeting other mitochondrial fission genes or other genes whose disruption promotes mitochondrial connectivity is sufficient to increase lifespan.

## Acknowledgments

Some strains were provided by the CGC, which is funded by NIH Office of Research Infrastructure Programs (P30 OD010440). We would also like to acknowledge the *C. elegans* knockout consortium and the National Bioresource Project of Japan for providing strains used in this research.

## Sources of Funding

This work was supported by the Canadian Institutes of Health Research (CIHR; http://www.cihr-irsc.gc.ca/; JVR; Application 399148 and 416150) and the Natural Sciences and Engineering Research Council of Canada (NSERC; https://www.nserc-crsng.gc.ca/index_eng.asp; JVR; Application RGPIN-2019-04302). JVR is the recipient of a Senior Research Scholar career award from the Fonds de Recherche du Québec Santé (FRQS) and Parkinson Quebec. The funders had no role in study design, data collection and analysis, decision to publish, or preparation of the manuscript.

## Disclosures

The authors have no conflicts of interest to declare.

## Author Contributions

Conceptualization: AT, JVR. Methodology: AT, JVR. Investigation: AT. Analysis: AT, JVR. Visualization AT, JVR. Writing – original draft: AT. Writing – review and editing: AT, JVR. Supervision: JVR.

## Materials & Correspondence

Correspondence and material requests should be addressed to Jeremy Van Raamsdonk.

## SUPPLEMENTAL FIGURES

**Figure S1.**
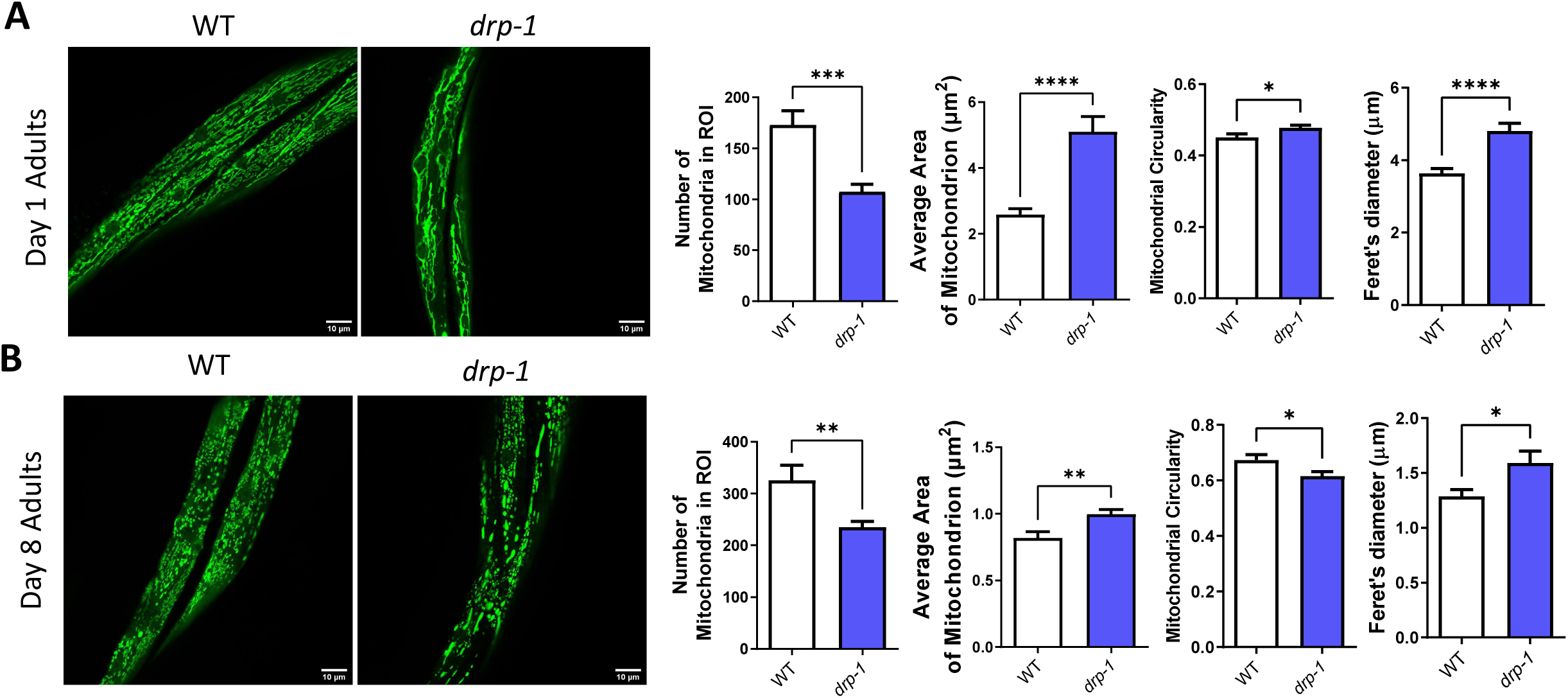
Disruption of *drp-1* increases mitochondrial network connectivity in wild-type animals. At day 1 of adulthood, disruption of *drp-1* in wild-type animals decreases the number of mitochondria, increases average mitochondrial area, increases circularity and increases ferret’s diameter (**A**). These measurements are similar at day 8, except that disruption of *drp-1* decreases circularity (**B**). Three biological replicates were imaged. Statistical significance was assessed using a student’s t-test. Error bars indicate SEM. *p<0.05, **p<0.01, ***p<0.001, ****p<0.0001. Scale bar indicates 10 µm.

**Figure S2.**
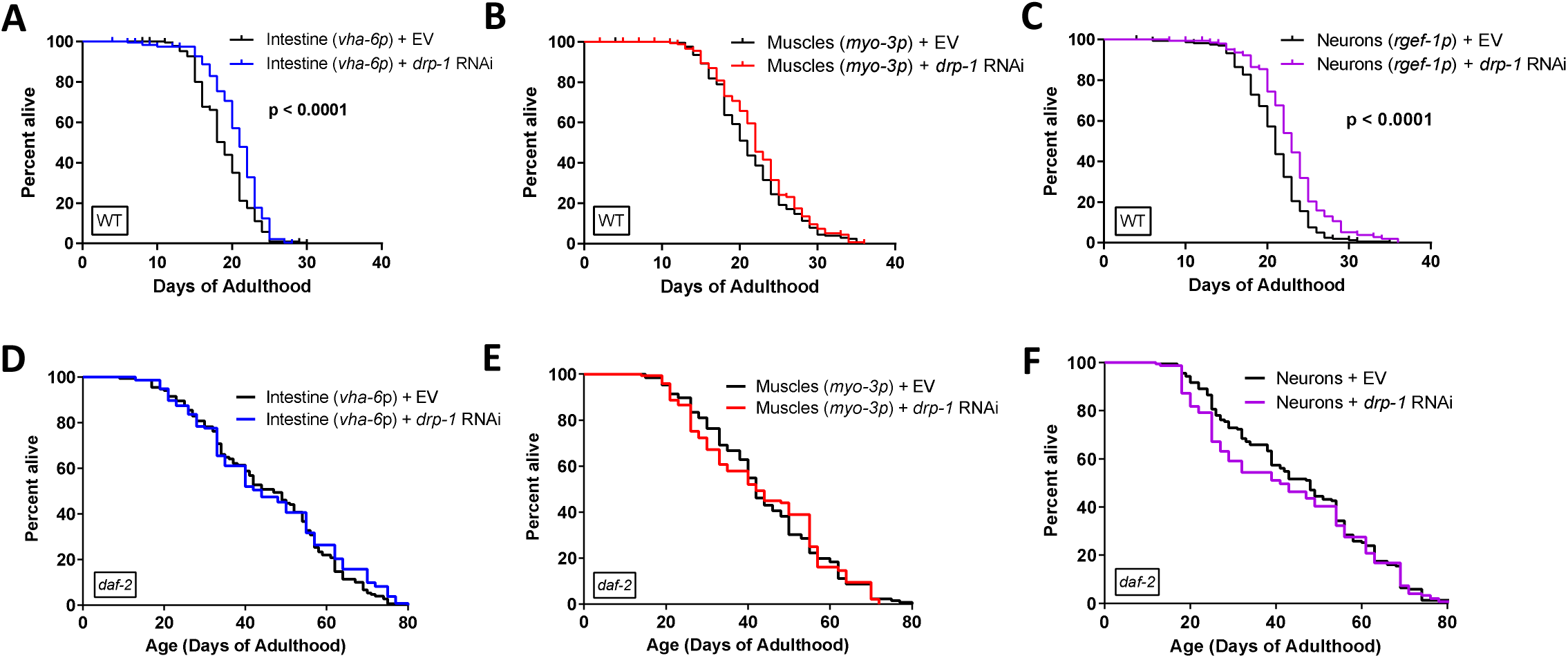
Tissue-specific knockdown of *drp-1* does not extend *daf-2* lifespan. Compared to animals fed empty-vector (EV) control bacteria, wild-type animals with disruption of *drp-1* in the intestine had increased lifespan Disruption of *drp-1* in the muscles of wild-type animals did not increase lifespan compared to animals fed EV Disruption of *drp-1* in the neurons of wild-type animals increases lifespan compared to animals fed EV (**C**). In *daf-2* animals, disruption of *drp-1* in the intestine, muscles or neurons of *daf-2* animals does not affect lifespan compared to animals fed EV (**D, E, F**). Four biological replicates were performed. Statistical significance was assessed using the log-rank test. *drp-1* RNAi data is from Figure 4 where control is worms with a *sid-1* mutation treated with *drp-1* RNAi.

**Figure S3.**
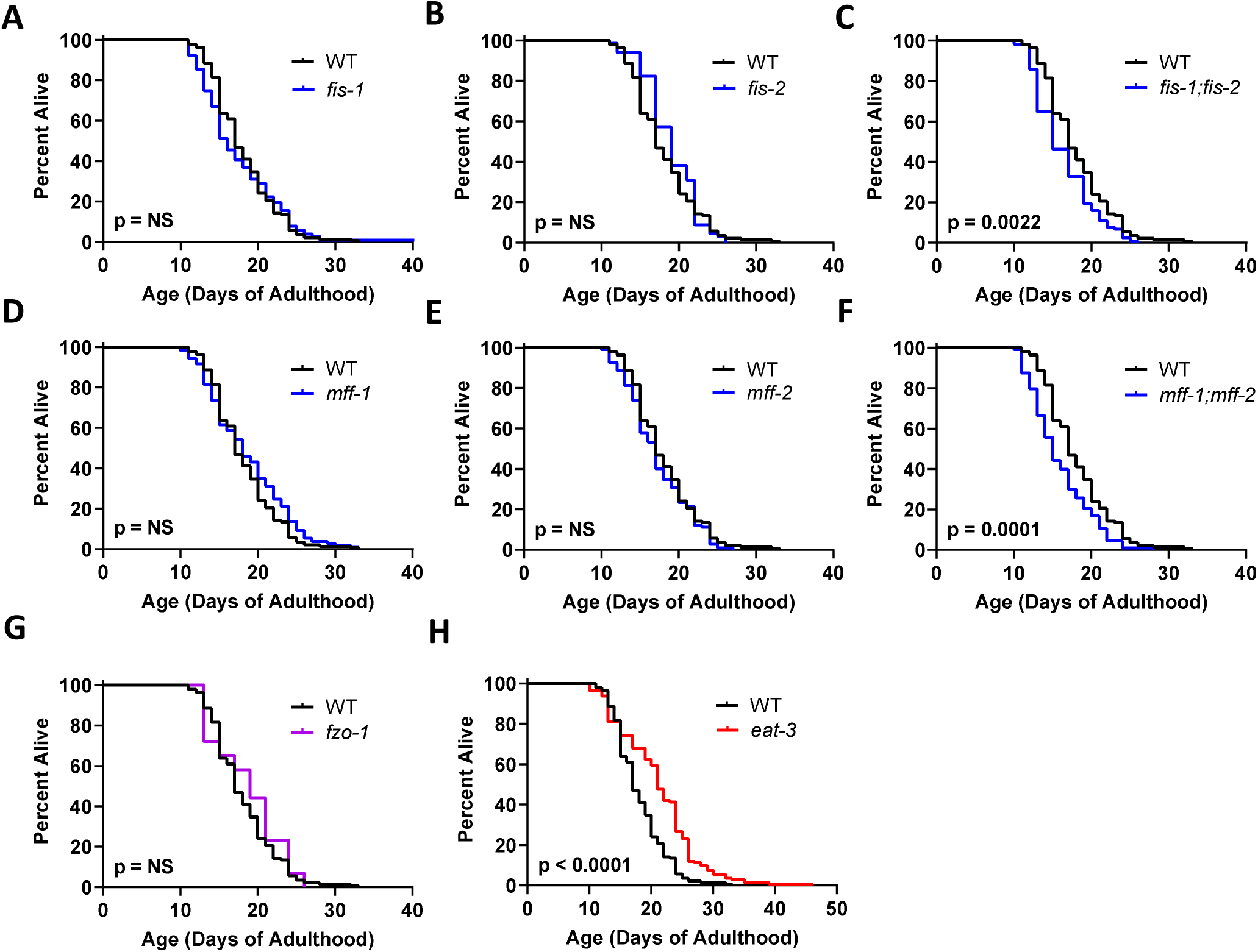
Effect of disrupting mitochondrial fission or fusion genes on lifespan in wild-type worms. Disruption of either *fis-1* (**A**), or *fis-2* (**B**) individually did not affect wild-type lifespan, while *fis-1;fis-2* double mutants exhibited a slight decrease in longevity (**C**). Similarly, *mff-1* (**D**) and *mff-2* (**E**) single mutants have a normal lifespan, while *mff-1;mff-2* double mutants exhibit decreased longevity (**F**). While deletion of the mitochondrial fusion gene *fzo-1* did not affect lifespan (**G**), disruption of *eat-3* extended longevity (**H**). At least three biological replicates were performed. Statistical significance was assessed using a log-rank test.

**Figure S4.**
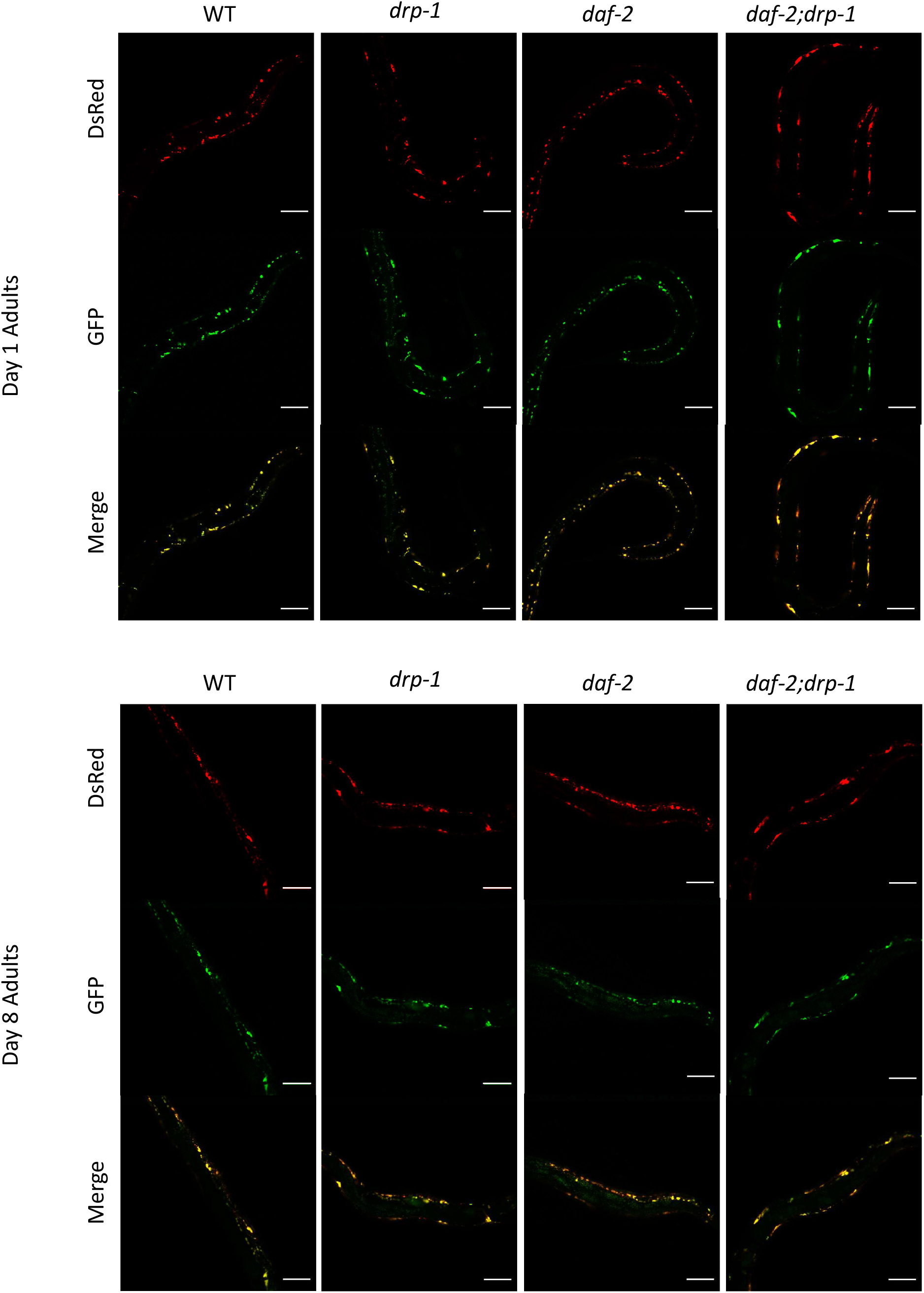
Visualization of the mtRosella mitophagy reporter reveals that disruption of *drp-1* increases mitophagy in *daf-2* worms in day 1 young adults. Whole body images of animals expressing the mtRosella mitophagy reporter were obtained using dual-channel confocal microscopy. Mitophagy levels were determined by comparing the fluorescence intensity of pH-insensitive DsRed to pH-sensitive GFP. See **Figure 8** for quantification of fluorescence. Representative images showing the compared intensity of each fluorophore were obtained by merging the two channels together. Scale bar represents 50 µm.

## Notes

### Competing Interest Statement

The authors have declared no competing interest.

